# Top-down Proteomics for the Characterization and Quantification of Calreticulin Arginylation

**DOI:** 10.1101/2024.08.08.607245

**Authors:** Richard M. Searfoss, Xingyu Liu, Benjamin A. Garcia, Zongtao Lin

**Affiliations:** Department of Biochemistry and Molecular Biophysics, Washington University in St. Louis, St. Louis, MO 63110

**Author notes:** These authors contributed equally (Richard M. Searfoss, Xingyu Liu). ***Corresponding Authors*** Benjamin A. Garcia - Department of Biochemistry and Molecular Biophysics, Washington University in St. Louis, St. Louis, Missouri 63110, United States;, Zongtao Lin - Department of Biochemistry and Molecular Biophysics, Washington University in St. Louis, St. Louis, Missouri 63110, United States.

## Abstract

Arginylation installed by arginyltransferase 1 (ATE1) features an addition of arginine (Arg) to the reactive amino acids (*e*.*g*., Glu and Asp) at the protein *N*-terminus or side chain. Systemic removal of arginylation after ATE1 knockout (KO) in mouse models resulted in heart defects leading to embryonic lethality. The biological importance of arginylation has motivated the discovery of arginylation sites on proteins using bottom-up approaches. While bottom-up proteomics is powerful in localizing peptide arginylation, it lacks the ability to quantify proteoforms at the protein level. Here we developed a top-down proteomics workflow for characterizing and quantifying calreticulin (CALR) arginylation. To generate fully arginylated CALR (R-CALR), we have inserted an R residue after the signaling peptide (AA1-17). Upon overexpression in ATE1 KO cells, CALR and R-CALR were purified by affinity purification and analyzed by LCMS in positive mode. Both proteoforms showed charge states ranging from 27-68 with charge 58 as the most intense charge state. Their MS2 spectra from electron-activated dissociation (EAD) showed preferential fragmentation at the protein *N*-terminals which yielded sufficient *c* ions facilitating precise localization of the arginylation sites. The calcium-binding domain (CBD) gave minimum characteristic ions possibly due to the abundant presence of >100 D and E residues. Ultraviolet photodissociation (UVPD) compared with EAD and ETD significantly improved the sequence coverage of CBD. This method can identify and quantify CALR arginylation at absence, endogenous (low), and high levels. To our knowledge, our work is the first application of top-down proteomics in characterizing post-translational arginylation *in vitro* and *in vivo*.

## Introduction

Post-translational modifications (PTMs) are important regulators of protein functions and cellular processes. Arginylation, one of the PTMs, is an essential component of the Arg/N-degron pathway which canonically targets the *N*-terminus of proteins leading to their destabilization and shortened half-lives in cells^1^. Arginylation is enzymatically generated by arginyl transferase 1 (ATE1) after adding arginyl residue to *N*-terminal Asp and Glu directly or Cys, Asn and Gln indirectly involving Cys oxidation and Asn/Gln deamidation^2^. The Arg/N-degron pathway also serves as a sensor for hypoxia and oxidative stress^1,3^, and plays a key role in autophagic protein degradation via lysosomes^4^. Other non-degradative roles of arginylation include protein secretion^5^ (*e*.*g*., serum albumin^6^), and intracellular translocation^1^ (*e*.*g*., β-actin^7^). Different from *N*-degron pathway, side chain arginylation was also observed in some proteins (*e*.*g*., α-synuclein^8^) and regulates protein folding and function. The possibility of arginylation on both *N*-terminus and side chain has significantly expanded the potential of arginylation in controlling a broad scope of protein substrates and their biological functions.

Several methods have been developed to identify the arginylation substrates systematically using bottom-up proteomics. The first proteomic analysis was carried out in 2006 on rat brain cytosol where six proteins were identified as ATE1 substrates^9^. In 2007, a global analysis of arginylation using antibody enrichment and proteomics identified 43 putative proteins with arginylation from mouse tissues^10^. Researching the same data against side chain arginylation further revealed 19 putative protein substrates^11^. Using the pan-arginylation antibody against side chain arginylation, 15 high-confidence proteins were identified as arginylation substrates^12^. Recently, our group has taken advantage of isotopic arginine labeling and developed the first unbiased proteomic platform, “activity-based arginylation profiling (ABAP)”, capable of identifying hundreds of *bona fide* arginylation sites from complex human proteomes^13^. These profiling methods have greatly expanded the understanding of the substrate scope of ATE1 in mammalian systems.

Among ATE1 substrates, calreticulin (CALR) was initially identified in rat brain with a subcellular translocation at the endoplasmic reticulum (ER)^9^. The primary function of CALR is controlling the newly synthesized proteins as a chaperone in the lumen of the ER. In addition, it is also a calcium (Ca^2+^) binding protein and is playing a pivotal role in Ca^2+^ homeostasis and signaling^14^. After ribosomal translation, CALR co-translationally cleaves the signaling peptide (AA 1-17) and translocates to the ER lumen as a mature protein^14^. The resulting *N*-terminal E18 site in CALR was arginylated by ATE1 leading to altered metabolic stability and half-life in the cytoplasm^14,15^. Although CALR arginylation (R-CALR) has been validated in the literature^14,15^, cellular levels of arginylation have not been evaluated due to the lack of analytical methods. In addition, the arginylation field has mainly focused on arginylation site discovery using bottom-up proteomics which is unable to quantify unmodified and arginylated proteins at the protein level. In contrast, top-down proteomics has arisen as a powerful approach for comprehensive analysis of proteoforms. We aim to take advantage of top-down proteomics for the identification and quantification of CALR and R-CALR.

In this study, a top-down proteomic approach has been developed for the analysis of wild-type and arginylated CALR. Both proteoforms have been produced and purified to compare their MS1 and MS2 behaviors. Several fragmentation methods have been used to compare fragmentation efficiencies and protein sequence coverage, the result showed EAD, ETD and UVPD all can characterize *N*-term arginylation with UVPD as a superior strategy for producing fragments from large proteins like CALR. Endogenous arginylation levels of CALR were identified by MS2 and quantified by MS1 spectra. Complementary to our previous bottom-up ABAP workflow, we showed that CALR were isotopically labeled by a novel on-bead ATE1 arginylation assay and quantified for arginylation levels thus served as proof-of concept data for “top-down ABAP”. Our work demonstrated an important step forward in arginylation studies using top-down proteomics; and provides an analytical approach for the identification and quantification of protein arginylation from biological systems.

## Experimental

### Plasmids

Plasmids encoding CALR and R-CALR were prepared by molecular cloning for mammalian expression. The fragment containing the coding sequence of human full-length CALR was from clone ID OHu23892 (GenScript Biotech, Piscataway, NJ, USA). The *C*-terminal version of the Halo tag was amplified from pFC14A (Promega, Madison, WI, USA). *C*-terminal Halo-tagged CALR was cloned into the pcDNA5/FRT vector (Invitrogen, Carlsbad, CA, USA). 18R-CALR was generated by inserting an Arg codon after the signal peptide (amino acids 1-17) of CALR using mutagenesis. The fragment containing the coding sequence of human full-length ATE1 was from clone ID OHu12263D (GenScript). *C*-terminal FLAG-tagged ATE1 was cloned into the pcDNA3.1+/C-(k)DYK vector (GenScript). Empty pcDNA3.1 plasmid (mock) was purchased from GenScript. Full sequences of plasmids are provided in the **Supporting Information**.

### Cell culture and transfection

HEK293T cells were purchased from ATCC. HEK293T ATE1 knockout (KO) cells were produced by CRISPR technology and validated in our previous study^13^. Cells were cultured in Dulbecco’s Modified Eagle’s Medium (DMEM) supplemented with high glucose, *L*-glutamine, and sodium pyruvate, along with 10% (*v*/*v*) fetal bovine serum (FBS) (R&D Systems, Minneapolis, MN, USA), at 37 °C in a humidified atmosphere containing 5% CO_2_^16,17^. For transfection, HEK293T ATE1 KO cells were plated in 100 mm tissue culture plates and allowed to attach and grow overnight. When cells reached approximately 70% confluency, each plate was transfected with a total of 2 µg of plasmids using FuGENE® HD (Promega). Briefly, plasmids and transfection reagent were premixed in 200 µL of OPTI-MEM I Reduced Serum Medium (Invitrogen), incubated at room temperature for 30 min, and then 800 µL of OPTI-MEM medium was added before dropwise addition to the cells. For plasmids expressing Halo-tagged CALR or 18R-CALR, 500 ng of the corresponding plasmid was added to each sample. The remaining 1.5 µg of co-transfected plasmids consisted of various ratios of ATE1 expression plasmids and empty pcDNA3.1, enabling different amounts of ATE1 co-expression in each sample. After 2 days of incubation, cells were harvested on ice and centrifuged. The resulting pellets were snap-frozen with liquid nitrogen and stored at -80 °C.

### Capture and Purifications of Halo-tagged CALR

Halo purification was performed according to the manual of HaloTag® Mammalian Pull-Down Systems (Promega). Briefly, thawed cell pellets were lysed with 300 µL Mammalian Lysis Buffer with the addition of benzonase (Sigma-Aldrich, Burlington, MA, USA) and protease inhibitors (Promega). The mixture was pressed through a 25-gauge needle 5 times for complete lysis. Crude lysates were centrifuged at 20,000 × g for 10 minutes at 4 °C, and supernatants (∼ 350 µL) were collected and added 200 µL TBS buffer. 500 µL of lysate was incubated with 50 µL Magne® HaloTag® Beads (Promega), the remaining lysate was saved as input sample for gel analysis. After rotating overnight at 4 °C, beads were washed with cold Wash Buffer 5 times (1 mL/wash), and 100 µL of 8 M urea in 50 mM TEAB buffer (pH 8.5) containing 2 mM TCEP. After 30 min incubation at room temperature, beads were washed with 200 µL of 50 mM TEAB buffer (pH 8.5) containing 2 mM TCEP and 2 mM CaCl^2^. Beads were then eluted with 50 µL of 50 mM TEAB buffer (pH 8.5) containing 2 mM TCEP and 5 U of AcTEV Protease (Invitrogen). After incubation at room temperature for 1 hour with shaking, supernatants were transferred to new tubes for mass spectrometry analysis.

### Detection of proteins with a Halo tag

For protein expression in the input lysate, 4 µL of each sample (0.8% of input) was analyzed by SDS-PAGE using a 4-12% NuPAGE™ Bis-Tris Gel in MOPS SDS Running Buffer (Invitrogen). For detection of bait expression, Halo-tagged proteins were labeled with HaloTag® TMR Ligand (Promega) in the lysate and imaged in gel (Excitation at 520 nm and Emission at 570-610 nm) using a LI-COR Odyssey® M Scanner (LI-COR Biosciences, Lincoln, NE, USA) controlled by LI-COR® Acquisition Software v1.0.0.55.

### Western Blotting

The protein gel after SDS-PAGE was subjected to Western Blot analysis for ATE1 expression. Proteins were transferred from the gel to a nitrocellulose membrane (0.45 µm, Bio-Rad, Hercules, CA, USA) in Tris/Glycine buffer with 20% methanol using a Trans-Blot® SD Semi-Dry Transfer Cell (Bio-Rad). The membrane was blocked for 1 hour at room temperature using Intercept® (TBS) Protein-Free Blocking Buffer (LI-COR Biosciences). Primary antibodies were diluted in the same blocking buffer with 0.2% of Tween-20. Both primary antibodies, mouse anti-FLAG M2 (1:2500 dilution, cat # F3165, Sigma-Aldrich) and rat anti-ATE1 clone 6F11 (1:1000 dilution, cat # MABS436, EMD Millipore), were mixed and incubated together with the membrane overnight at 4 °C. Following primary antibody incubation, the membrane was washed 3 times for 5 minutes each with Tris-buffered saline containing 0.1% Tween 20 (TBST), then incubated with secondary antibodies for 30 minutes at room temperature. For detection of both the FLAG tag and ATE1, IRDye® 680RD Goat Anti-Mouse IgG and IRDye® 800CW Goat Anti-Rat IgG secondary antibodies (LI-COR Biosciences) were diluted in blocking buffer at 1:10,000 with 0.1% Tween-20. The membrane was then washed twice for 5 minutes with TBST and rinsed with TBS before being imaged. The FLAG tag and ATE1 signals on the same membrane were scanned in two different channels (700 channel for FLAG and 800 for ATE1) using the LI-COR Odyssey® M Scanner (LI-COR Biosciences).

### Silver staining

For silver staining analysis of purified CALR proteins eluted by AcTEV protease, 10% of the total elution from each sample was loaded and separated by SDS-PAGE using a 4-12% NuPAGE™ Bis-Tris Gel in MOPS SDS Running Buffer (Invitrogen). The gel was subsequently stained with the Pierce™ Silver Stain Kit (Thermo Fisher Scientific, Waltham, MA, USA) and imaged using the LI-COR Odyssey® M Scanner (LI-COR Biosciences) in the 470trans channel.

### On-bead ATE1 assay for arginylation of overexpressed CALR

Halo-tagged CALR was purified the same as described earlier but without AcTEV cleavage. Two CALR samples were prepared for isotopic labeling. 50 µL magnetic beads containing CALR were washed with 100 µL 1x buffer diluted from 5x reaction buffer (50 mM HEPES, 30 mM KCl, and 10 mM MgCl_2_). The ATE1 assay was re-constituted as an on-bead arginylation assay based on previous studies^13,18^. Assay components were added individually to beads on ice. A 20-μL reaction was set up containing 1x reaction buffer, 2 mM Arg (Arg^0^ and Arg^10^, respectively), 2 mM ATP, 3 μM tRNA^Arg^, 1 μM RARS1, 3 μM human ATE1, and beads as substrate. The assay was incubated at 37 °C for 1 h with beads disturbed. Beads were then washed to remove assay components by 100 µL 1x buffer, followed by 200 µL of 50 mM TEAB buffer (pH 8.5) containing 2 mM TCEP and 2 mM CaCl_2_. Beads were then eluted with 50 µL of 50 mM TEAB buffer (pH 8.5) containing 2 mM TCEP and 5 U of AcTEV Protease (Invitrogen). After incubation at room temperature for 1 hour with shaking, supernatants were transferred to new tubes for mass spectrometry analysis. 5 µL of elution was used for silver staining.

| ATE1 assay composition |  |  |  |  |  |
| --- | --- | --- | --- | --- | --- |
| order | reagent | stock concentration | volume (μL) | volume (μL) | final concentration |
| 1 | ddH<sub>2</sub>O | N.A. | 7 | 7 |  |
| 2 | Reaction buffer | 5x | 4 | 4 | 1x |
| 3 | Arg | 40 mM | 1 (Arg<sup>0</sup>) | 1 (Arg<sup>10</sup>) | 2 mM |
| 4 | ATP | 10 mM | 4 | 4 | 2 mM |
| 5 | CALR | On beads | NA | NA | NA |
| 6 | tRNA | 60 μM | 1 | 1 | 3 μM |
| 7 | RARS1 | 20 μM | 1 | 1 | 1 μM |
| 8 | ATE1 | 30 μM | 2 | 2 | 3 μM |

### ATE1 *in vitro* assay for arginylation of commercial CALR

Commercially available CALR with a His tag and an open 18E as *N*-terminal was purchased from Abcam (catalog No.: ab276554) and dissolved in 1x reaction buffer to a concentration of 0.5 µg/μL. 1x buffer was diluted from 5x reaction buffer (50 mM HEPES, 30 mM KCl, and 10 mM MgCl_2_). The arginylation assay was set up on ice by mixing a 20-μL reaction containing 1x reaction buffer, 2 mM Arg, 2 mM ATP, 3 μM tRNA^Arg^, 1 μM RARS1, 3 μM human ATE1, and 3.5 µg CALR protein as substrate. The reconstituted assay was incubated at 37 °C for 1 h. The CALR was diluted to 100 ng/µL and stored at -80 °C before LCMS analysis.

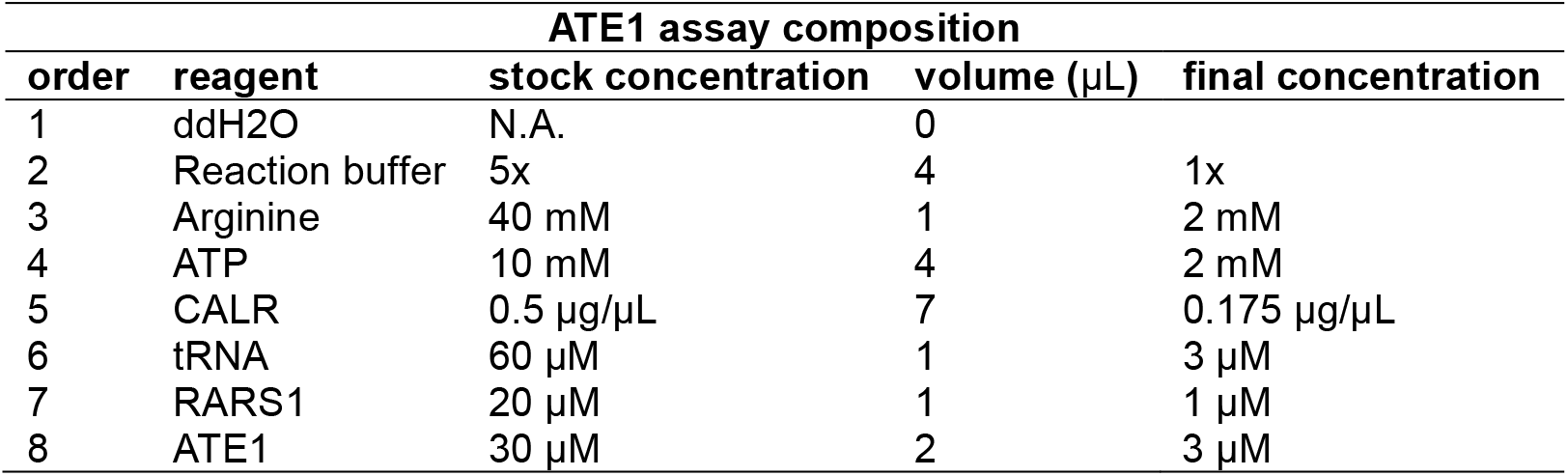

### LCMS analysis of CALR proteins using ZenoTOF 7600

An LC-MS/MS system consisting of a Waters Acquity UPLC M-Class or an Eksigent M5 MicroLC System coupled to a Sciex ZenoTOF 7600 in MRM^HR^ mode was used for intact protein analysis. Protein samples were maintained at 7 °C on sample tray in LC. Separation of proteins was carried out on a Waters nanoEase M/Z Protein BEH C4 Column (1.7 µm, 300 Å, 300 µm X 50 mm) at 50 °C. Solvents consisted of 0.1% formic acid in water and 0.1% formic acid in acetonitrile for buffers A and B, respectively. A flowrate of 10 µL/min was used to apply a linear gradient at 3% B for 1 minute, increased from 3% to 20% B in 2 minutes, from 20% to 40% B in 12 minutes, from 40% to 90% B in 1 minute, held at 90% B for 1 minute, from 90% to 3% B in 1 minute, and held at 3% B for 2 minutes. The ZenoTOF 7600 was operated with the Optiflow TurboV microflow ion source applying a spray voltage of 4300 V. MS1 scans were conducted over a scan range of *m/z* 500-2000 with a 0.1 sec accumulation time. MS2 scans were conducted with MRM^HR^ selecting 3 charge states from each target protein with Q1 resolution set to “Low” and a 0.25 sec accumulation time. The scan range was set to *m/z* 100-3000 for initial experiments, but since no meaningful fragments were identified beyond *m/z* 2000, this was set as the upper limit for all later experiments. EAD fragmentation was conducted for 5 ms at 5000 nA beam current and 1 keV electron energy, with a declustering potential of 80 V and collision energy of 12 V.

### LCMS analysis of CALR proteins using Ascend

An LC-MS/MS system consisted of a Vanquish Neo UHPLC coupled to an Orbitrap Ascend (Thermo Scientific) was used for ETD and UVPD analysis. Proteins were maintained at 7 °C on the sample tray in LC. Separation of proteins was carried out on a Waters nanoEase M/Z Protein BEH C4 Column (1.7 µm, 300 Å, 150 µm X 100 mm) at 40 °C. Solvents consisted of 0.1% formic acid in water and 0.1% formic acid in acetonitrile for buffers A and B, respectively. A flowrate of 0.5 µL/min was used to apply a linear gradient at 2% B for 1 minute, increased from 3% to 30% B in 2 minutes, from 30% to 60% B in 6 minutes, from 60% to 90% B in 1 minute, held at 90% B for 1 minute, from 90% to 2% B in 1 minute, and held at 3% B for 3 minutes. The Thermo Fisher Ascend was operated in the Intact Protein mode with the Low-Pressure mode selected and a 2500 V spray voltage was applied to a 50 Å, 10-cm pulled fused silica emitter tip on the ESI source. MS1 scans were conducted over a scan range of *m/z* 700-2000 in both the ion trap and the orbitrap. The enhanced scan rate was selected in the ion trap with a standard AGC target and a maximum injection time of 200 ms. In the orbitrap, MS1 scans were conducted with a 120K resolution with 200% AGC target and 200 ms maximum injection time. MS2 scans were collected using a user-defined DIA window in center mass DIA window mode. UVPD MS2 scans were conducted in the orbitrap with 60K resolution from *m/z* 100-4000 with a 300% AGC target and 200 ms maximum injection time, and a 5 ms ETD reaction time and 1E5 ETD reagent target.

### Data analysis for top-down proteomics

All MS2 files (.WIFF and .RAW) were converted to .MZML using MSConvert ver. 3.0.23073 with the peak picking option selected^19^. The resulting .MZML files were then deconvoluted using FLASHDeconv hosted under OpenMS v3.1.0, with “write detail” option set to true and “threads” set to 4^20^. All other parameters were left with default settings. Visualization of deconvoluted MS2 data was performed in ProsightLite v1.4 using monoisotopic precursor mass and M (neutral) fragment mass mode, with fragment matching set to 10ppm error, and either the “BY and CZ*” setting for EAD and ETD analyses or “UVPD 9” selected for UVPD analyses^21^. Further plotting of chromatograms and both MS1 and MS2 spectra was performed in RStudio (v2023.03.0+386).

## Results and Discussion

### Preparation of CALR and R-CALR

CALR resides in the ER where the signal peptide containing amino acids 1-17 is cleaved to release the mature CALR. Taking advantage of this process, we constructed a plasmid system to express mature CALR and R-CALR proteins adapted from a previous strategy^15^. To produce arginylated CALR, an amino acid R was added after the signal peptide which left a *N*-terminal R on CALR after maturation (**Fig. 1a**). CALR and R-CALR were overexpressed in HEK293T cells with ATE1 KO to ensure that CALR with an open *N*-terminal E17 was not arginylated by ATE1. The proteins were designed to possess a *C*-terminal Halo tag for covalent capture by HaloTag beads during affinity purification, allowing for the 8 M urea wash procedure to remove interacting proteins. TEV site was inserted between CALR and Halo tag and was cleaved by TEV protease to yield purified CALR and R-CALR proteins ready for mass spectrometry analysis (**Fig. 1**). To monitor the protein overexpression, cell lysates were separated by gel electrophoresis and treated with Halo-TMR reagent. The Halo signals showed that both CALR-Halo species were overexpressed in HEK293T cells with and without ATE1 co-overexpression (**Fig. 1b**). Western blotting using anti-flag and anti-ATE1 antibodies confirmed the absence of endogenous expression of ATE1 in HEK293T cells. After TEV cleavage, the purified CALR proteins were visualized after silver staining which showed at expected molecular weights without Halo tag.

**Figure 1.**
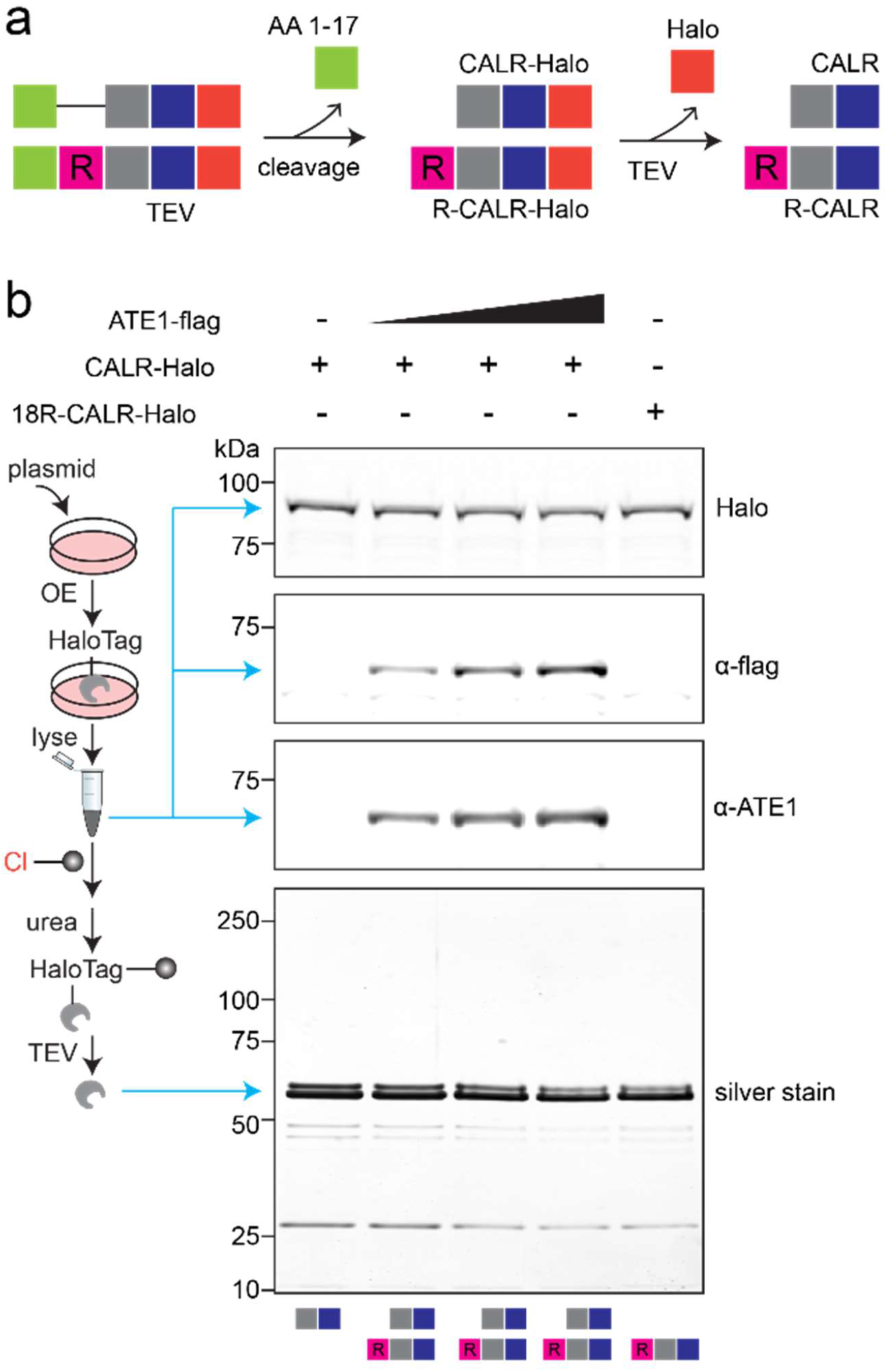
Affinity pulldown and purification of CALR and R-CALR proteins. **a**, schematic for design and purification strategies of CALR proteins. The signal peptide was co-translationally cleaved, and the resulting mature proteins were bound to HaloTag beads. Upon removal of interacting proteins by 8 M urea, proteins were cleaved from beads by TEV protease to yield the desired proteins. **b**, gel visualization of overexpression and purified CALR proteins. Proteins with and without Halo tag were shown with different molecular weights on gels. Anti-flag and anti-ATE1 antibodies were used for western blotting against the ATE1 protein.

### Configuration and Design of LC-MS Analysis Method

The top-down platform was first started using the Sciex ZenoTOF 7600 coupled with a microflow LC system (**Fig. 2a**). Using this platform, we were able to apply microflow chromatographic LC separation to the analysis of intact proteins, providing reproducible and robust protein retention and elution. Following others in the top-down field^22^, a C4 reversed-phase column was used for less protein retention compared with C18, allowing for easier elution and less carryover from run to run. We also opted for a 300 Å, 300 µm X 50 mm pore and particle size in a shorter length column to prevent clogging of the column with a larger (∼47 kDa) calreticulin protein as well as to keep back pressure lower while running a high flow rate. The above components allowed for consistent chromatographic peaks around 30 seconds in width (**Figure S1**). For mass spectrometry analysis, we chose electron-based fragmentation because collision-induced dissociation has historically provided insufficient fragmentation of intact proteins. EAD fragmentation has proved useful in other experiments for intact proteins, and there is a plethora of literature supporting ETD and UVPD. Typically, the larger the protein, the less electron KE is needed because the protein is more positively charged to pull in electrons. Lower reaction time can sometimes facilitate obtaining sufficient sequence coverage. However, in the case of calreticulin, a slightly longer reaction time was still needed due to the chemical nature of the protein. This method provided the required sequence coverage of the *N*- and *C*-terminus of calreticulin, thus allowing for experimentation in this study.

**Figure 2.**
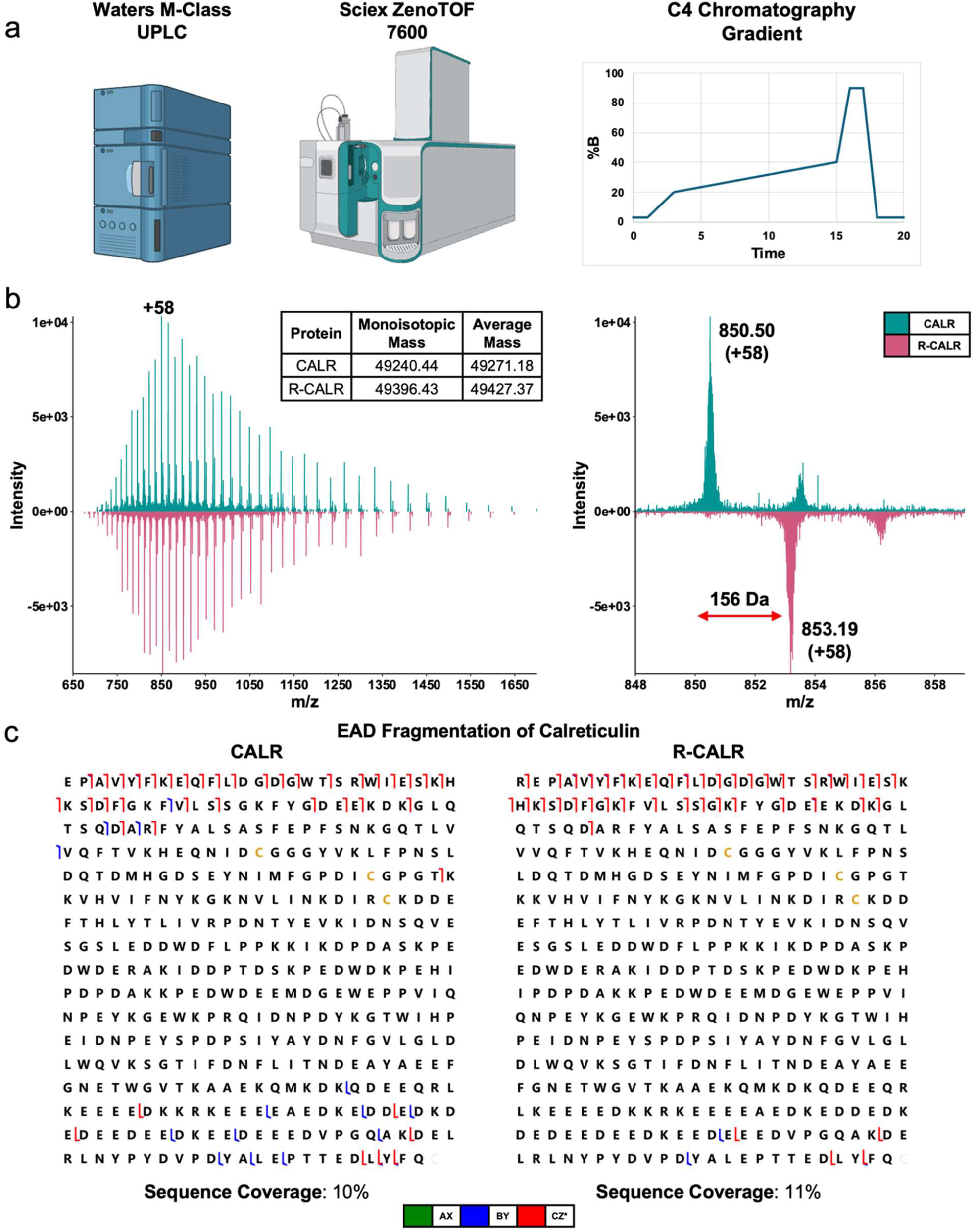
MS1 and MS2 characterization of CALR and R-CALR proteins. **a**, Top-down proteomics workflow used for characterizing intact calreticulin using a C4-reversed phase separation chromatographic gradient. **b**, MS1 charge state distribution and measured monoisotopic and average masses of CALR and R-CALR shown as an average scan across the chromatographic peak. Visualization of the MS1 mass shift incurred by adding an arginine residue to the *N*-terminus, focusing on charge state +58. **c**, MS2 sequence coverage of CALR and R-CALR when subjected to EAD fragmentation.

### Identification of Wild-Type and Arginylated Calreticulin with LC-MS and EAD Fragmentation

To begin developing a top-down proteomics method for identifying and quantifying arginylated calreticulin, we first sought to classify the protein ensuring adequate fragmentation and identification of the shift in precursor and fragment masses due to arginylation. Using the purified CALR and R-CALR from lane 1 and lane 5 in **Fig. 1b**, the proteins were run individually and subjected to either EAD fragmentation on the Sciex 7600 ZenoTOF, or ETD or UVPD fragmentation on the Ascend. MS1 level analysis collected on the Sciex 7600 shows a strong signal from CALR, observing charge states +27 to +68 (**Fig. 2b**), and a clear shift of 156 Da between the CALR and R-CALR species (**Fig. 2b**). The observed monoisotopic and average masses of these species following deconvolution also agrees with the predicted masses calculated within a 10-ppm error. The results demonstrated that differentiation of the arginylated form of calreticulin is possible. For MS2 analyses on the Sciex 7600 ZenoTOF, three charge states for each species were targeted using an MRM^HR^ analysis: [M+60H]^60+^ (*m/z* 822.18 for CALR, *m/z* 824.78 for R-CALR), [M+54H]^54+^ (*m/z* 913.42 for CALR, *m/z* 916.31 for R-CALR), and [M+50H]^50+^ (*m/z* 986.41 for CALR, *m/z* 989.54 for R-CALR). EAD fragmentation yielded a 10% sequence coverage of CALR and an 11% sequence coverage of R-CALR (**Fig. 2c, Figure S2**).

It is challenging to use the electron-based fragmentation method to analyze the fragmentation of this protein efficiently. This is largely due to both its large size (∼47 kDa) and the high abundance of aspartic acid and glutamic acid residues surrounding its calcium-binding domain (Uniprot ID: P27797), accounting for almost 27% of the total residues of this protein (114/423 AA’s, **Figure S3**). This results in a high density of negative charge in this region, which is less suitable for electron fragmentation. However, we still got full sequence coverage of the *N*-terminus of the protein in both unmodified and modified species, allowing us to identify the absence and presence of the arginylation modification. When comparing the MS2 EAD spectra of CALR versus R-CALR (**Figure S2**), a strong signal from the *c*_*1*_ ion (*m/z* 174.13) of the *N*-terminal R-CALR can be seen as expected, owing to the high fragmentation probability of a positively charged arginine residue. No signal can be seen from the CALR *c*_*1*_ ion (*m/z* 147.08) because the *N*-terminal aspartic acid residue is followed by a proline, which is poor at fragmenting by electron-based methods^23,24^. However, a series of signals from the *c*_*2*_, *c*_*3*_, and *c*_*4*_ ions are also identified, thus validating the absence of the *N*-terminal R in the CALR sample.

### Identification of Wild-Type from Arginylated Calreticulin with ETD and UVPD Fragmentation

We then sought to characterize these proteins with ETD fragmentation, which relies on a different electron delivery mechanism than EAD, and UVPD which performs backbone fragmentation with photons instead of electrons. MS1 analysis of CALR and R-CALR showed similar charge state distributions as the Sciex ZenoTOF 7600 (**Figure S4a**). This is beneficial because it shows that similar analysis can be performed on TOF and Orbitrap instruments, as well as utilizing microflow and nano-flow systems, and they can be expected to generate consistent results for this intact protein. For MS2 analyses on the Ascend, three charge states were targeted using a fixed-window DIA analysis across both fragmentation experiments: [M+58H]^58+^ (*m/z* 850.57 for CALR, *m/z* 853.25 for R-CALR), [M+55H]^55+^ (*m/z* 896.81 for CALR, *m/z* 899.66 for R-CALR), and [M+51H]^51+^ (*m/z* 967.06 for CALR, *m/z* 970.11 for R-CALR). ETD fragmentation produced a 25% sequence coverage of CALR and a 26% sequence coverage of R-CALR, showing full sequence coverage of the *N*-terminus and allowing differentiation of the modification from wild-type (**Fig. 3, Figure S4b**,**c**). UVPD fragmentation was performed over triplicate runs and scans averaged together to boost sequence coverage. This is common in top-down proteomics because UVPD often causes internal fragmentation of proteins due to multiple pulses of the UV laser, as well as results in several types of fragments being formed^25,26^. However, UVPD is still beneficial because it provides the highest sequence coverage of proteins across fragmentation types when applied to intact proteins^27-30^. In this experiment using ProsightLite to visualize data with UVPD 9 option selected, 9 total ion types were mapped (*a, a*+, *x, x*+, *b, y, y*-, *c*, and *z*)^22,31-33^. As a result, we achieved 74% and 61% sequence coverage of CALR and R-CALR, respectively, providing the highest sequence coverage of this protein and its calcium-binding domain (**Fig. 3, Figure S4b**,**d**). Together, each method is amenable to identifying calreticulin in future experiments.

**Figure 3.**
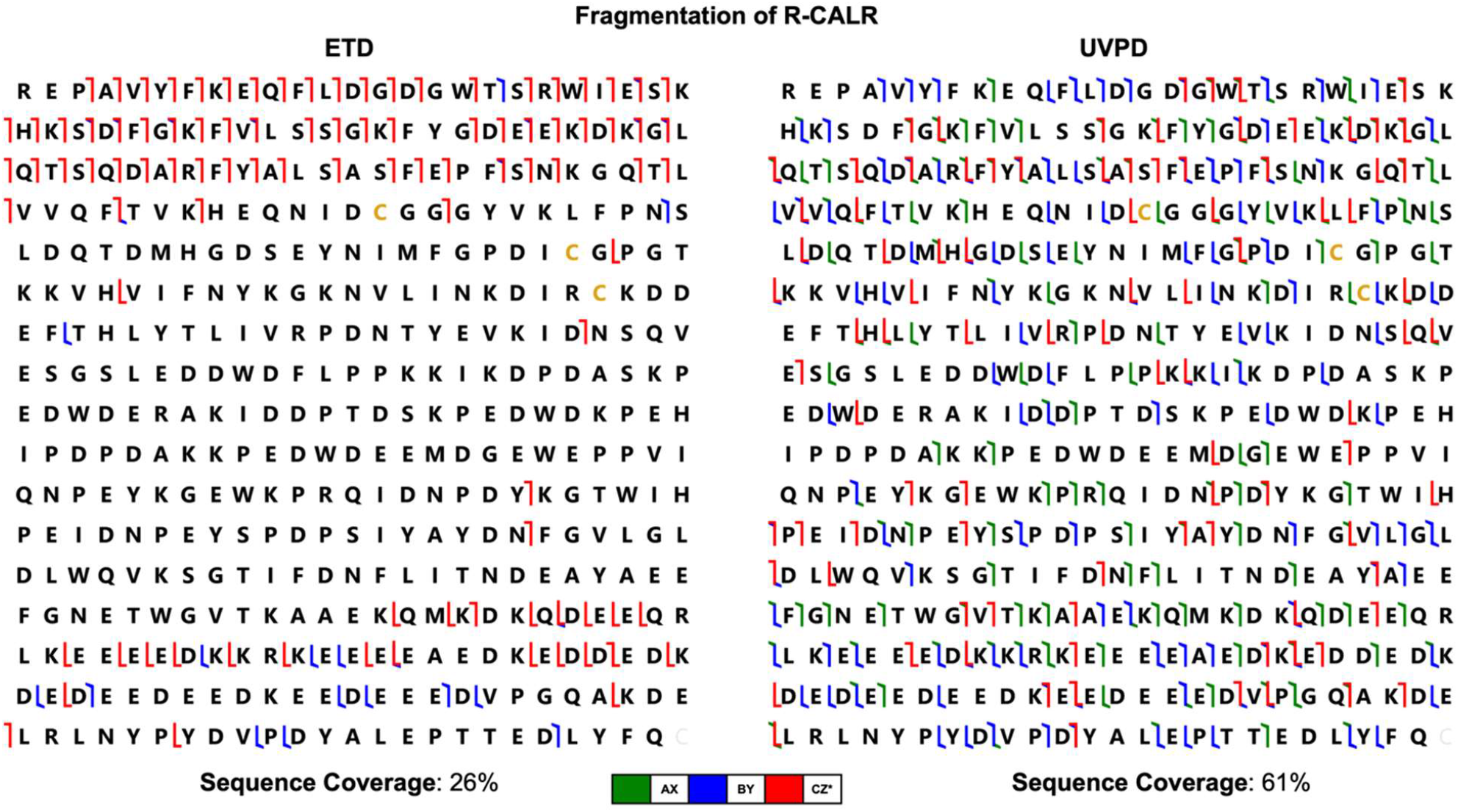
MS2 characterization of R-CALR protein using ETD and UVPD fragmentation. Both approaches still allow visualization of the *N*-terminus of the protein with localization of the modification using the *c*-ion series. UVPD further provides protein-wide sequence coverage and nine types of produced ions.

### WT-CALR and R-CALR Mixing for Proteoform Quantitation

With confident fragmentation and coverage of R-CALR and identification of specific *c*-ions localizing the arginyl modification, we next tested the quantitation of co-occurring CALR species. There is no consensus in the top-down proteomics field on the most correct or accurate way to quantify an intact proteoform, but multiple approaches have been proposed. Label-free, metabolic labeling and chemical labeling are the three primary approaches for quantifying proteoforms. However, the precise mechanism of quantitation from the spectral level still varies, whether that be at a single charge state’s extracted ion chromatogram (XIC) peak area, an XIC of multiple charge states, or the deconvolved level, and most studies have focused on the smaller intact mass range <30 kDa^34-36^. As such, we show multiple strategies, each exemplifying the ability to recapitulate the mixed quantities of CALR and R-CALR. CALR and R-CALR showed comparable expression levels before and after purification (**Fig. 1**), so we moved forward with this mixing experiment assuming an equal amount (1:1) of both species. Six total mixtures of each species were created by volume alone, here denoted as WT for CALR and R for R-CALR. We sequentially increased the concentration of WT while decreasing the concentration of R, with final WT: R ratios of 2:8, 4:6, 5:5, 6:4, 8:2, and 10:1. We first performed an extracted ion chromatogram (XIC) of the [M+58H]^58+^ peaks, corresponding to *m/z* 850.50 and *m/z* 853.20 for WT and R respectively, of 0.2 Da window (± 0.1 Da). Each XIC was Gaussian smoothed with a 5-point smoothing window in the SciexOS software to reduce the effect of signal fluctuation. A clear increase in peak area and intensity can be seen in the WT from the lowest 2:8 ratio to the highest 10:1 ratio of WT as compared to the steady decrease in peak area and maximum intensity from the highest 2:8 ratio to the lowest 10:1 ratio of R (**Fig. 4a**). The percent signal of WT and R was calculated by dividing peak area of WT or R by the sum peak areas (WT + R). This measurement showed an increase from 12% WT in the 2:8 ratio to 82% WT in the 10:1 ratio, demonstrating the quantitative abilities of the top-down proteomics platform for co-eluting proteoforms (**Figure S5**).

**Figure 4.**
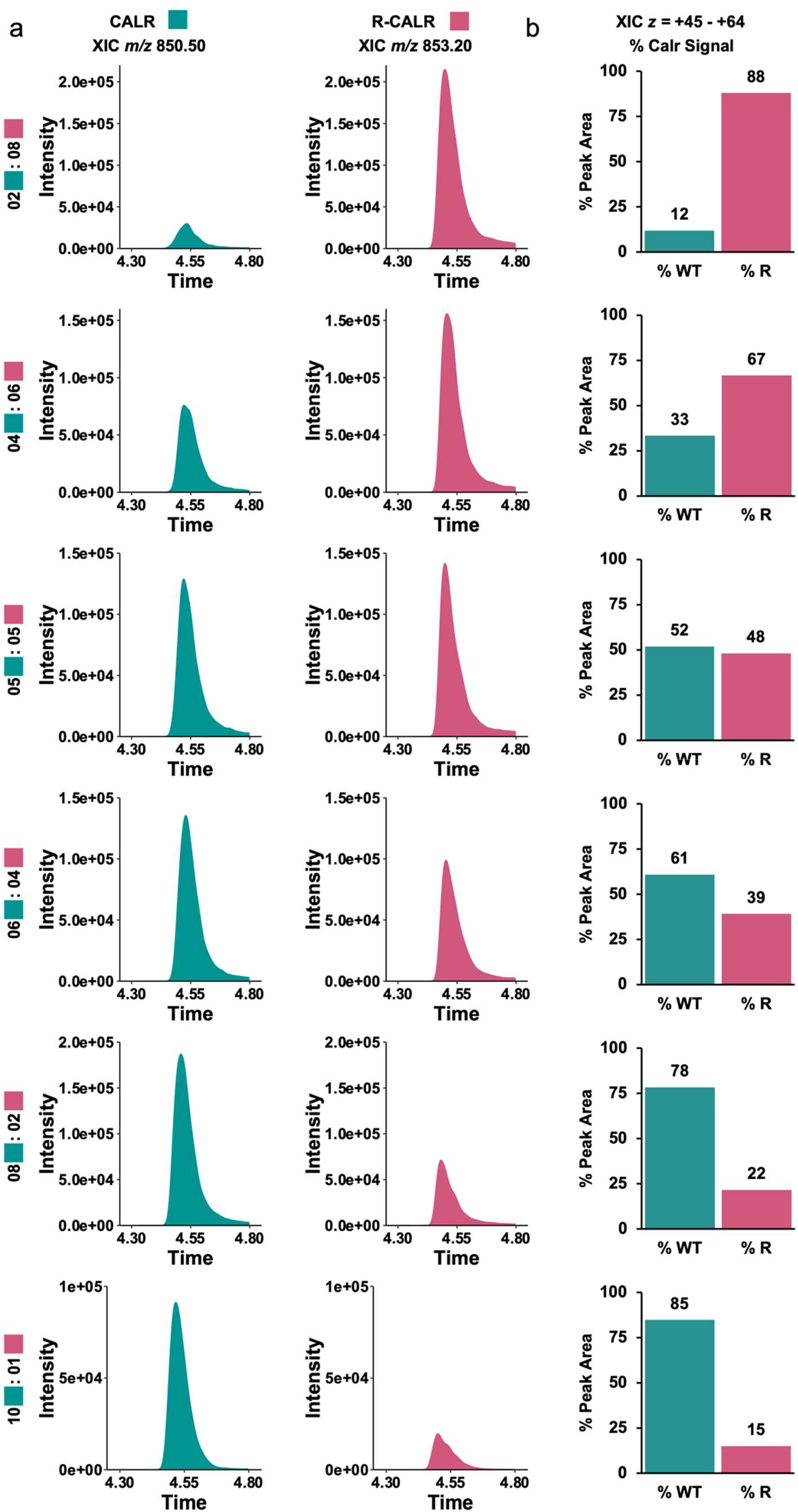
MS1 Label-free quantitation of CALR and R-CALR in mixed populations. **a**, XIC of the [M+58H]^58+^ charge state of unmodified CALR, representing an increasing quantity in each sequential experiment, and XIC of the [M+58H]^58+^ charge state of modified R-CALR, representing a decreasing quantity in each sequential experiment. **b**, The percent of the total calreticulin signal measured in each experiment represented by either CALR or R-CALR. The total calreticulin signal was calculated by XIC of twenty charge states of each species and summing their chromatographic peak areas.

Given that individual charge states could have variable ionization efficiencies, especially considering the addition of a positively charged arginyl modification, we also sought to find a more averaged value for each species^37,38^. We performed another XIC for each species, covering 20 total charge states from [M+45H]^45+^ to [M+64H]^64+^ (**Figure S6**), and each XIC was gaussian smoothed in SciexOS with 5 point smoothing window. The peak area for each charge state was summed to get a total signal for the WT and R species in each mixture, and the percent signal of WT and R was calculated by dividing the sum peak area of WT or R by the total sum intensity (WT + R). A steady increase is seen in the relative signal of WT and a decrease in the relative signal of R, with the measured 4:6, 5:5, 6:4, 8:2 ratios of WT:R almost exactly matching the theoretical ratios (**Fig. 4b**). We also wanted to visualize the signal intensity changes within the raw data. The average MS1 spectra were created by averaging all MS1 scans within the chromatographic window corresponding to the CALR species. Taking a focused look into the [M+64H]^64+^ peaks, corresponding to *m/z* 770.86 for WT and *m/*z 773.30 for R, we observed the intensities of each species increasing and decreasing relative to each other, following the pattern of how WT and R were mixed (**Figure S7**). These approaches demonstrate the ability of our platform to quantify arginylated calreticulin in the presence of the wild-type form, thus supporting the prospect of identification and quantitation in biological samples.

### Identification and Quantification of Arginylated Calreticulin from Cells

To begin probing the ability to identify and quantify endogenous arginylation, CALR was overexpressed in ATE1 KO cells with an open E18 residue at the *N*-terminus, and sequentially co-overexpressing ATE1 enzyme to install the *N*-terminal arginyl modification (**Fig. 1b**). R-CALR was used as a positive control for quantification. These five samples represent different levels of arginylation: absence (lane 1, **Fig. 1b**), endogenously low (lanes 2-4, **Fig. 1b**), and high (lane 5, **Fig. 1b**). To quantify the unmodified and modified calreticulin species, an XIC of 0.2 Da window (± 0.1 Da) and Gaussian smoothing with a 2-point smoothing window was performed on [M+61H]^61+^ (*m/z* 808.73 for CALR, *m/z* 811.29 for R-CALR), [M+57H]^57+^ (*m/z* 865.41 for CALR, *m/z* 868.15 for R-CALR), and [M+54H]^54+^ (*m/z* 913.43 for CALR, *m/z* 916.32 for R-CALR) in the SciexOS software. The resulting peak areas were summed and the percent arginylated was found. While R-CALR showed MS1 spectra for arginylation, CALR (with or without ATE1 overexpression) did not show clear MS1 signals at individual charge states for R-CALR due to no or low (endogenous) levels of arginylation. To confirm the absence or presence of arginylation, an MRM^HR^ was performed targeting the same three R-CALR charge states. The resulting EAD MS2 spectra did not observe any signal from the N-terminal *c*_*3*_ ion (*m/z* 400.23) from sample 1, thus confirming the absence of R-CALR species in CALR sample without ATE1 expression (**Fig. 5a, Figure S8a**).

**Figure 5.**
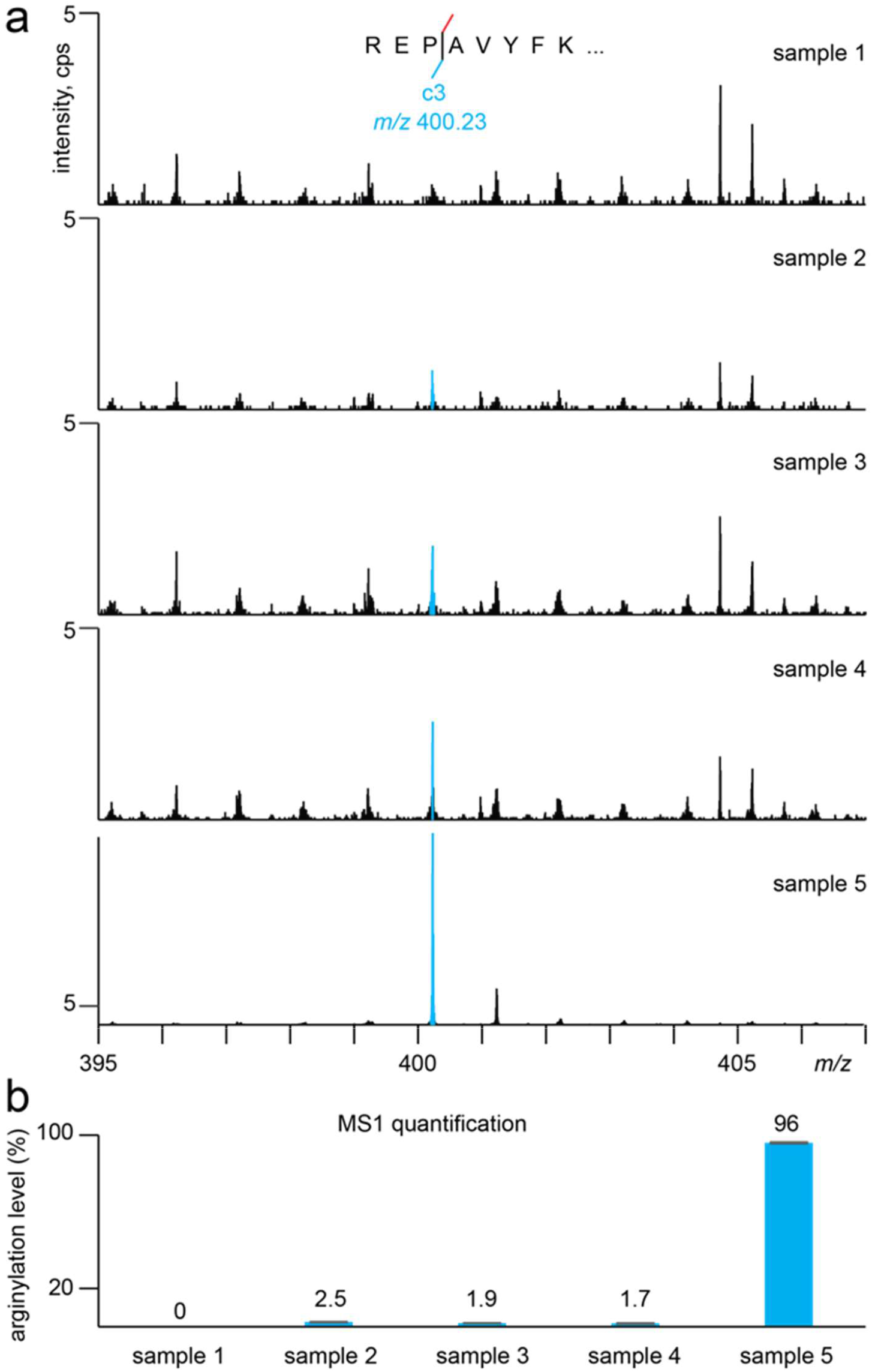
Identification of endogenously arginylated calreticulin in the presence of co-overexpressed ATE1 enzyme. **a**, MS2 spectra confirming the absence of *N*-terminal arginylation in the control sample, as indicated by the lack of the *c*_*3*_ ion (*m/z* 400.23), and confirming the presence of *N*-terminal arginylation, indicated by the low-level presence of the *c*_*1*_ ion after transient expression of ATE1 enzyme in samples 2-4 and high signal in sample 5. **b**, The percent signal of R-CALR from the total MS1 calreticulin signal, calculated from XIC of three charge states of each species.

Using this information, we similarly wanted to confirm the presence of R-CALR with EAD MS2 fragmentation to validate our quantitation. Samples 2, 3, and 4 all had overexpression of calreticulin with transient co-expression of ATE1 enabling possible installation of low-level *N*-terminal arginylation. Applying the same MRM^HR^ method with EAD fragmentation, the presence of the low-level *N*-terminal *c*_*3*_ ion (*m/z* 400.23) is easily observable across samples 2-4, thus providing confident data for the presence of R-CALR proteoform (**Fig. 5a, Figure S8b-d**). Each sample showed endogenously low-level arginylation, ranging from 1.7-2.5% R-CALR which is approaching the MS1 limit of quantification (LOQ) for this instrument (**Fig. 5b**). Although we were unable to show clear MS1 signals for the R-CALR species for endogenous arginylation, our approach demonstrated the capability of using *c*-ion series in MS2 spectra for arginylation identification and using summed MS1 signals from three charge states (61+, 57+, 54+) to quantify endogenous arginylation.

Sample 5 as an arginylation control sample employed the arginyl modification in the protein sequence (**Fig. 1b**). The MRM^HR^ with EAD fragmentation showed a strong signal of the *N*-terminal *c*_*3*_ ion (*m/z* 400.23), further validating the findings of this ion in the previous samples 2, 3, and 4, and lack of this ion in sample 1 (**Fig. 5a, Figure S8e**). The resulting XIC of CALR and R-CALR in this sample showed 96% arginylation level (**Fig. 5b**) with a clear presence of CALR in MS1 spectra (**Figure S9a-c**). Although this sample should have provided a 100% R-CALR signal given genetic arginylation in the plasmid, it is possible that CALR may be formed as a product in cells. One possibility is that endogenous aminopeptidases such as AMPB (Uniprot ID: Q9H4A4) in cells may de-arginylate R-CALR to CALR even though this pathway has not yet been established in the arginylation field. Collectively, these findings provided the necessary evidence to support a promising approach for finding endogenously arginylated proteins at or near the LOQ of the instrument. This method showed the co-elution of CALR and R-CALR through multiple experiments. Such chromatographic behavior enabled arginylation identification at the MS2 level disregarding the intensities of their MS1 signals.

### Identification of Arginylated CALR from *in vitro* ATE1 Assay

With previous success in developing a bottom-up proteomics assay to detect new arginylation targets of ATE1 through *in vitro* and *ex vivo* labeling, we sought to test the application of top-down proteomics for arginylation site elucidation, a complementary approach to bottom-up ABAP^13^. We tested the suitability of our top-down method for detecting arginylated species using a commercial CALR (different sequence from overexpressed CALR in this study) and a routine in-solution ATE1 assay. EAD fragmentation targeting [M+63H]^63+^ (*m/z* 753.71), [M+58H]^58+^ (*m/z* 818.60), and [M+54H]^54+^ (*m/z* 879.17) confirmed the sequence of the protein, providing 28% sequence coverage and high fragmentation coverage of the *N*- and *C*-terminus (**Figure S10**). The deconvoluted monoisotopic mass was measured to be 47,394.45 Da, agreeing with the theoretical mass of 47,394.15 Da within a 10-ppm error. The results validate the identity of the commercial calreticulin and allow us to use this protein for the *in vitro* arginylation assay. Next, the CALR was incubated with ATE1 in solution to allow for the *N*-terminal installation of the arginyl modification (**Figure 6**). MS1 spectra clearly showed the R-CALR as a product with a similar charge state distribution pattern as the CALR. At charge state [M+56H]^+56^, we are still able to measure the mass shift incurred by the addition of the arginine, increasing from *m/z* 847.84 in CALR to *m/z* 850.63 in R-CALR (**Figure 6b,c**). Further, the observed deconvoluted monoisotopic mass of 47,550.26 Da was still within a 10-ppm error of the predicted monoisotopic mass of 47,550.25 for R-CALR. Here, we were able to demonstrate the ability of the top-down proteomics platform to identify the *in vitro* installation of an arginyl to CALR, both supporting the previously developed bottom-up proteomics assay and providing the framework to identify arginylation sites in pure proteins or biological proteomes.

**Figure 6.**
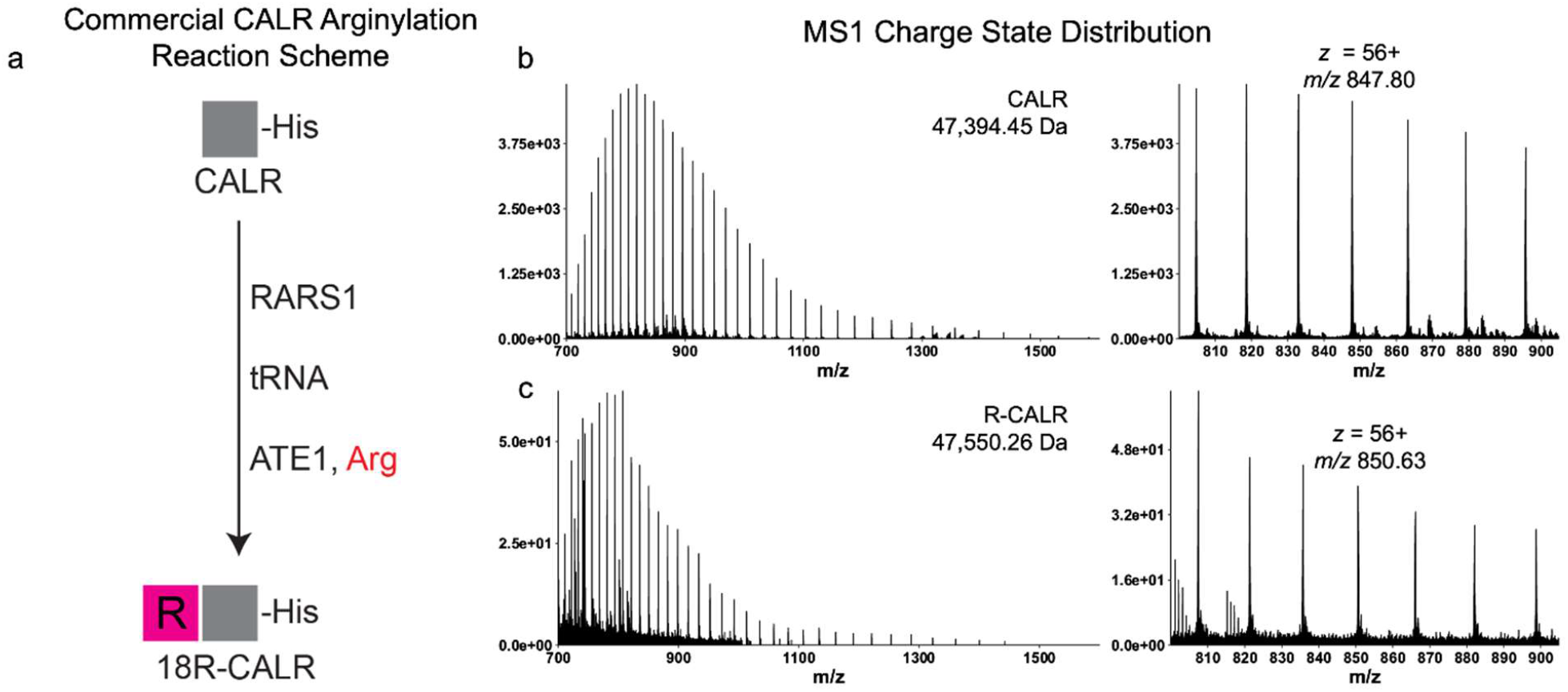
Proof of concept of in vitro arginylation assay for identification of endogenous arginylation events. **a**, Reaction scheme of RARS1 charging the R-tRNA, followed by incubation with ATE1 and CALR to allow *N*-terminal installation of the arginyl modification. **b**, MS1 charge state distribution and measured monoisotopic mass of commercial CALR shown as an average scan across the chromatographic peak, and a focused view of *m/z* 800-900 to visualize unmodified CALR peaks. c, MS1 charge state distribution and measured monoisotopic mass of commercial CALR after ATE1 installation of arginyl modification, shown as an average scan across the chromatographic peak. The focused view of *m/z* 800-900 allows visualization of mass shift compared to the unmodified CALR peaks.

### Identification and Quantification of arginylated CALR from On-bead *in vitro* ATE1 Assay

Moving one step forward from in-solution ATE1 assay, we arginylated CALR on beads by the ATE1 arginylation assay adapted from our previous work^13^. Briefly, RARS1, tRNA, ATE1 were added to beads with light or heavy arginine (R^0^ or R^10^, respectively) to arginylate CALR (**Fig. 7a**). The assay components were washed away from beads which were then eluted by AcTEV protease for CALR species. A silver stain showed the presence of the CALR band without the obvious presence of RARS1 and ATE1 from the assay (**Fig. 7a**). They were analyzed by top-down proteomics on the Sciex ZenoTOF 7600. EAD fragmentation targeting [M+60H]^60+^, [M+56H]^56+^, and [M+54H]^54+^ for both the R^0^ and R^10^ arginylated calreticulin allowed confident identification of the *N*-terminal arginyl modification, providing 15% and 17% sequence coverage of each species, respectively (**Figure S11a**). At the MS1 level, a clear shift in mass between the CALR and R-CALR can be seen in both samples and further, a slight shift can be seen between the R^0^-CALR and R^10^-CALR (**Fig 7b,c, Figure S11b, Figure S12**). In the MS2 spectra, both light and heavy R-CALR show the same fragmentation pattern and provide full coverage of the *N*-terminus of the protein (**Fig 7d,e, Figure S12**). Here, it is much clearer to see the shift in mass of 10 Da from the *c*-ions in the R^0^ versus the R^10^ fragment ions, showing consistent relative intensities to each other as well. To quantify the arginylation levels from the ATE1 assay, we extracted XIC peak areas of intensities of 3 charge states. The results showed that the arginylation efficiencies (60.3% for R^0^-CALR and 73.8% for R^10^-CALR) are higher than 60% in both samples (**Fig 7b,c**), with error bars showing the standard deviations of relative intensities of CALR to R-CALR at three charge states. These results are comparable with our previous data using the bottom-up approach showing a 64.7% arginylation efficiency by spectra count^13^, and confirm the success of isotopic top-down proteomics for monitoring *in vitro* arginylation, thus making it possible for unbiased elucidation of arginylation sites in other protein substrates of ATE1.

**Figure 7.**
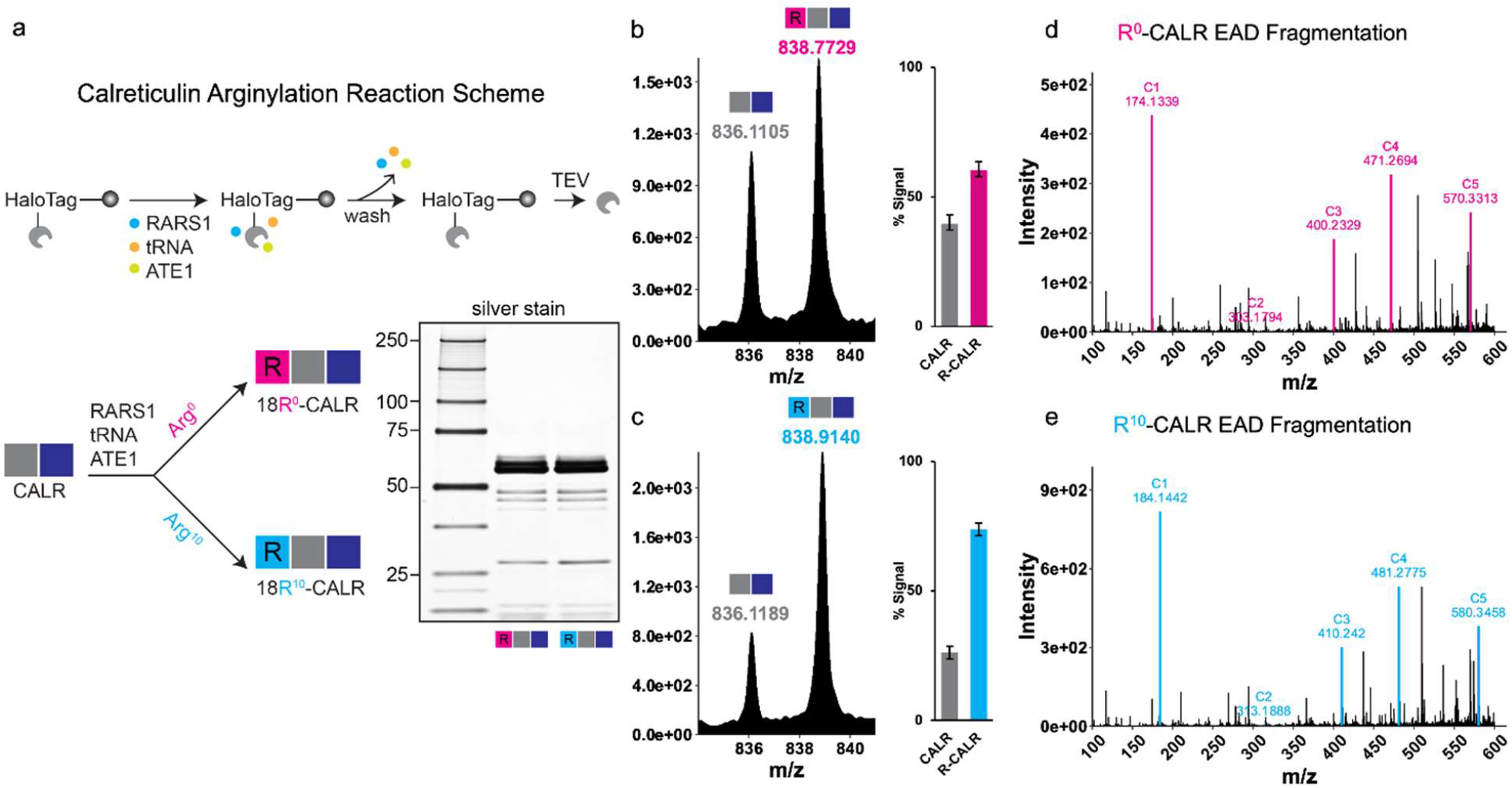
Proof of concept of *in vitro* arginylation assay for identification of endogenous arginylation events. **a**, Calreticulin arginylation scheme using Halo-tag purified calreticulin and light or heavy arginine. **b**, MS1 charge state *z* = 59 shows mass shift of CALR to R^0^-CALR, and XIC of three charge states shows arginylation efficiency of 60.3%. **c**, MS1 charge state *z* = 59 shows mass shift of CALR to R^10^-CALR, and XIC of three charge states shows arginylation efficiency of 73.8%. **d**, MS2 spectra confirms *N*-terminal installation of R^0^ modification with signature *c*_*1*_ ion (*m/z* 174.13). **e**, MS2 spectra confirms *N*-terminal installation of R^10^ modification with signature *c*_*1*_ ion (*m/z* 184.14), and a shift of 10 Da of all *c*-ions from the R^0^ spectra shown above.

## Conclusions

There is a lack of methods to characterize post-translational arginylation in proteins using a top-down proteomics approach. In this study, we purified unmodified and arginylated CALR at various levels using affinity purification. The MS1 and MS2 spectra were obtained and compared to reveal the differences between CALR and R-CALR. *N*-terminal arginylation was identified by a series of signature c ions after EAD fragmentation. EAD fragmentation increased the protein sequence coverage up to 17% in canonical calreticulin, and increased to 28% in the commercial calreticulin, primarily calculated from *N*-terminal and *C*-terminal fragments while the other fragments covering the internal sequence were not frequently obtained. Similar results were observed using ETD fragmentation, demonstrating the consistency and comparability between the two methods. Comparing the EAD, the UVPD fragmentation has boosted the sequence coverage up to 61%, potentially facilitating the MS2 characterization of big proteins such as CALR. When mixing CALR and R-CALR proteins at different ratios, our method can efficiently identify and quantify the amounts of two co-eluting protein species. Additionally, we demonstrated the capability of this workflow in the confident identification and quantification of endogenous arginylation levels, even at endogenously low levels pushing the limits of quantitation of our platform. To complement our previous bottom-up approach using ABAP, we show that isotopic arginylation can be analyzed by top-down proteomics. The established workflow can characterize and quantify the arginylated CALR, thus paving the way for studying the arginylation of other ATE1 substrate proteins from the human proteome. Since arginylation is involved in various biological processes, this top-down method has great potential to facilitate revealing the cellular functions of arginylated proteins.

## Supporting Information

The Supporting Information is available free of charge at https://pubs.acs.org/doi/.

- MS1 and MS2 spectra of CALR species.
- Detailed plasmid and protein sequences.

## Author Information

### Author Contributions

B.A.G. and Z.L. conceived the project. R.M.S. and X.L. designed and performed the experiments and prepared figures. Z.L., R.M.S., and X.L. drafted the manuscript. R.M.S., X.L., B.A.G., and Z.L. analyzed data, edited, and approved the manuscript.

### Notes

Z.L. and B.A.G. are co-founders of LasNova Therapeutics, LLC.

## Acknowledgments

B.A.G. thanks partial support from NIH NS111997, NIH HD106051, and NSF CHE 2127882 grants. Z.L. thanks partial support from WUSM BMB Seed Grant PJ000027587 and Research Education Component through NIA P30 AG066444 grant.

