## Supporting Information for "Top-down Proteomics for the Characterization and Quantification of Calreticulin Arginylation"

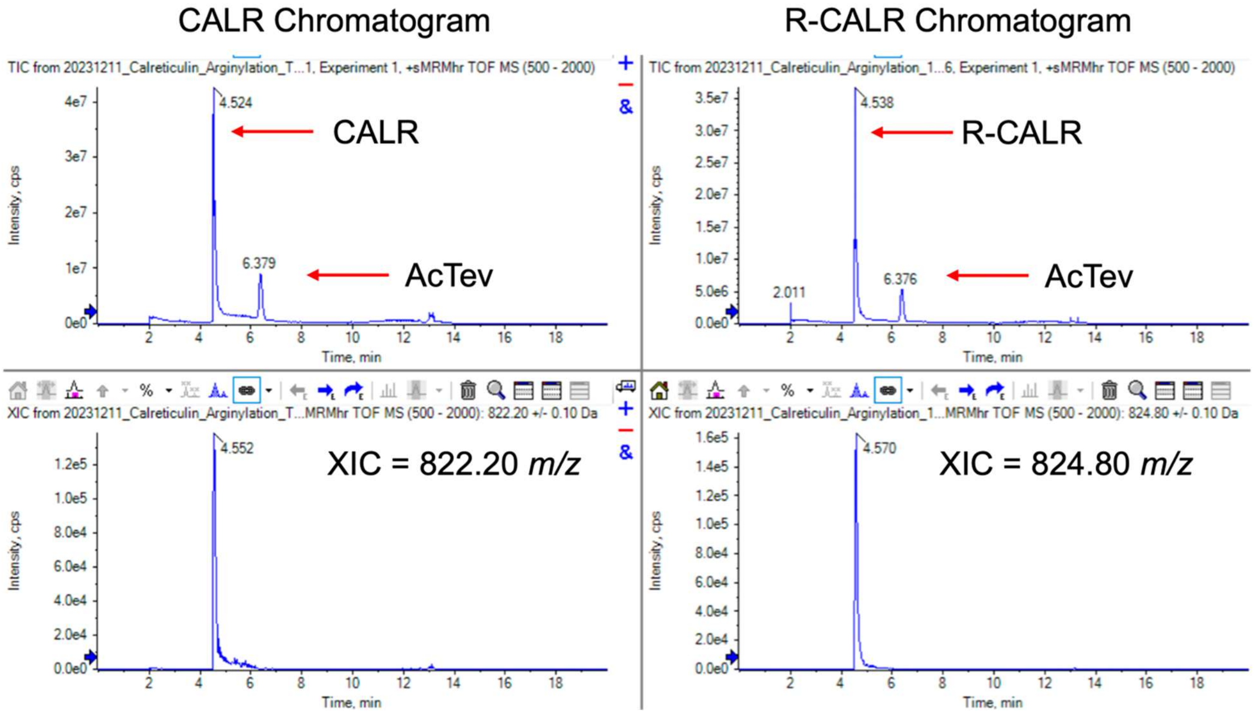


**Supplemental Figure 1.** Chromatogram of CALR and R-CALR, and XIC of a single charge state of each species, showing tight chromatographic peaks of roughly 30 seconds.

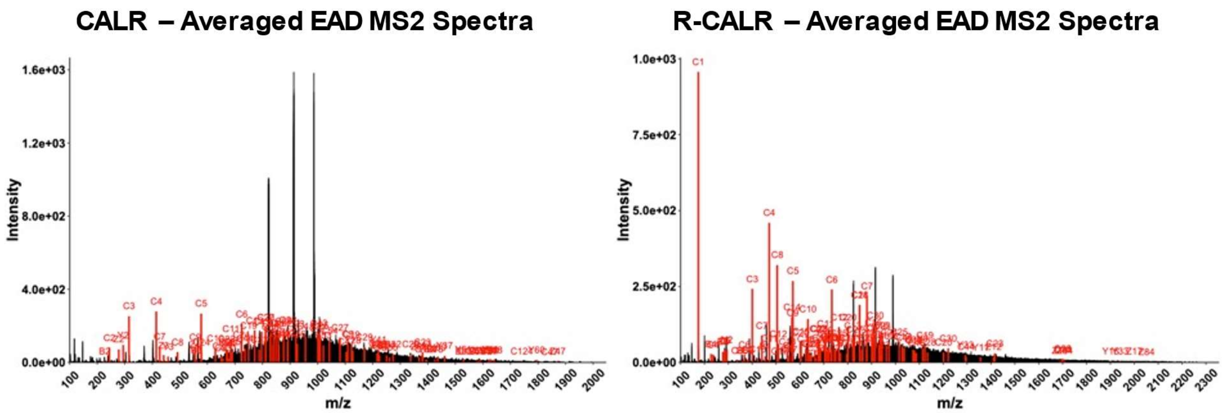


**Supplemental Figure 2.** MS2 spectra of CALR and R-CALR EAD fragmentation. Clear difference is observed between the *c*-ion patterns, with the *c1*-ion (*m/z* 174.13), representing the fragmentation of the N-terminal R residue, is clearly visible in the R-CALR MS2 spectrum.

A.

| Calreticulin Composition | | |
|---|---|---|
| **AA** | **Count** | **Percent** |
| A | 15 | 3.55 |
| C | 3 | 0.71 |
| D | 58 | 13.71 |
| E | 56 | 13.24 |
| F | 19 | 4.49 |
| G | 25 | 5.91 |
| H | 7 | 1.65 |
| I | 19 | 4.49 |
| K | 42 | 9.93 |
| L | 21 | 4.96 |
| M | 4 | 0.95 |
| N | 17 | 4.02 |
| P | 29 | 6.86 |
| Q | 16 | 3.78 |
| R | 9 | 2.13 |
| S | 19 | 4.49 |
| T | 17 | 4.02 |
| V | 18 | 4.26 |
| W | 11 | 2.60 |
| Y | 18 | 4.26 |

B.

C.

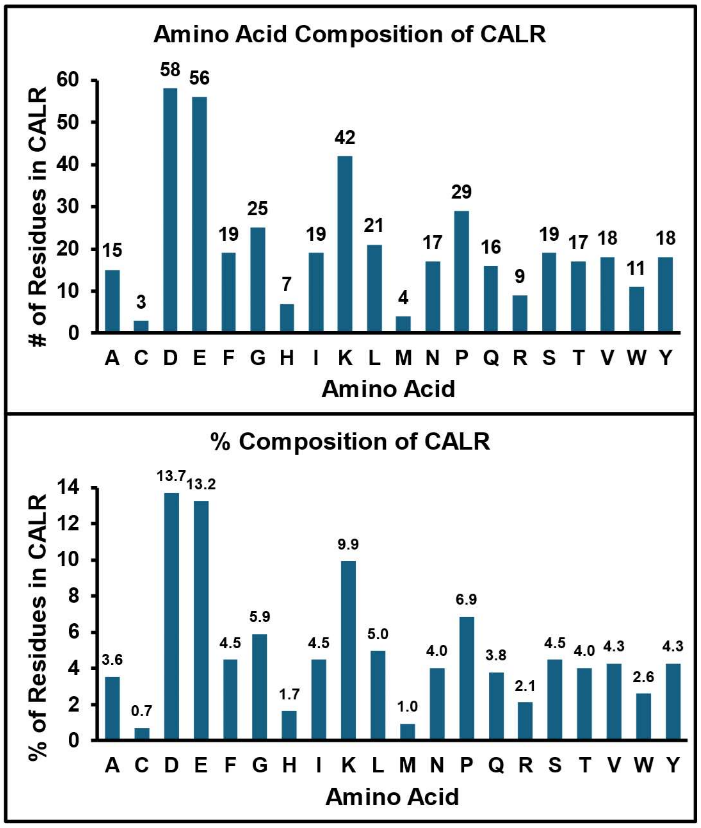


**Supplemental Figure 3.** Amino acid composition of calreticulin. **a,** Table depicted the number of each amino acid comprising calreticulin, and the percent that each amino acid represents in the full-length protein. **b,** Visualization in bar graph of the amino acid composition by total number of each residue in calreticulin. **c,** Visualization in bar graph of the amino acid composition by percent of each residue in calreticulin.

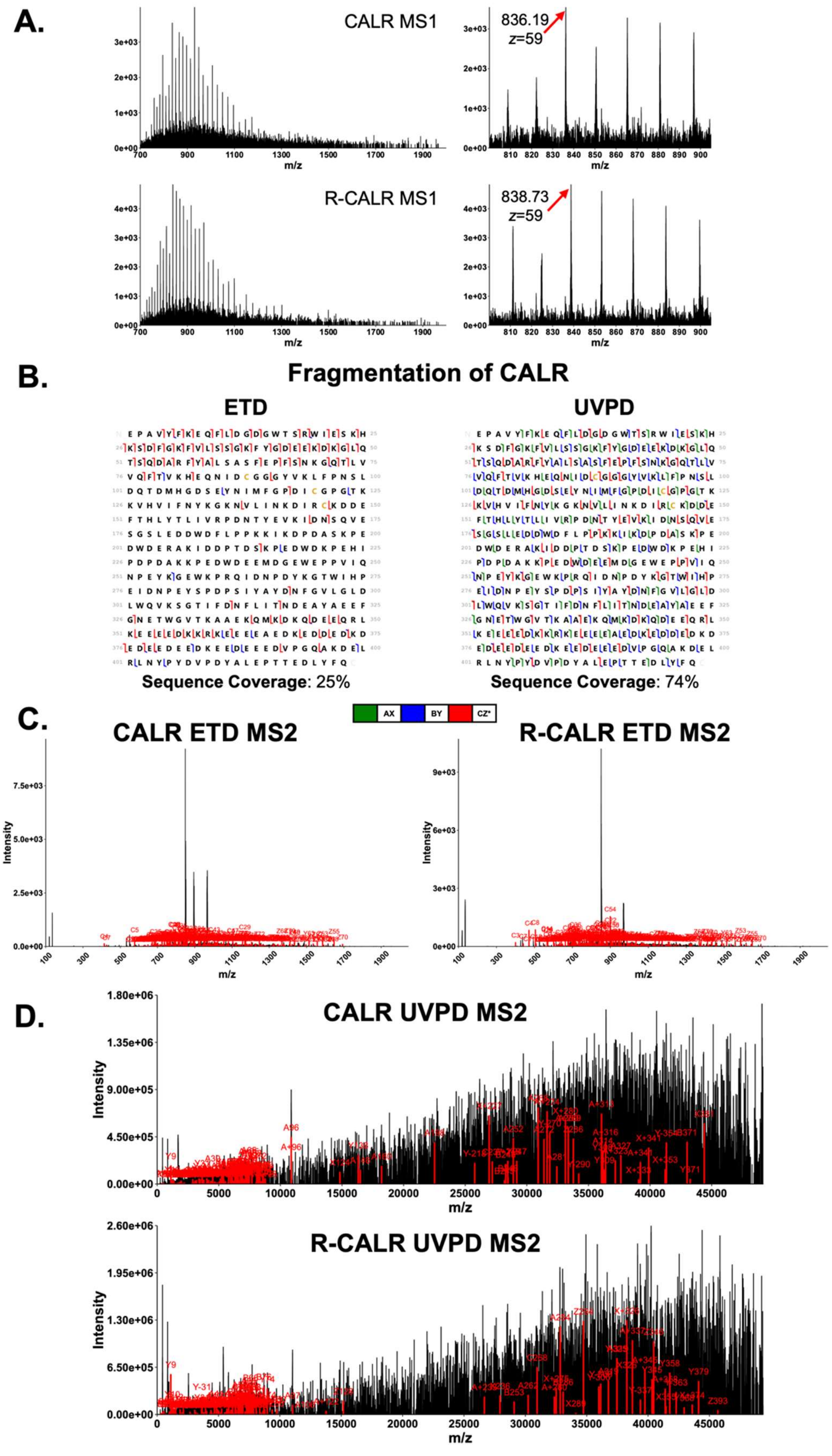

A.
CALR MS1
836.19
z=59
R-CALR MS1
838.73
z=59
m/z
B.
Fragmentation of CALR
ETD
UVPD
Sequence Coverage: 25%
Sequence Coverage: 74%
AX
BY
CZ*
C.
CALR ETD MS2
R-CALR ETD MS2
Intensity
m/z
D.
CALR UVPD MS2
R-CALR UVPD MS2
Intensity
m/z

**Supplemental Figure 4.** Fragmentation of CALR with ETD and UVPD fragmentation on the ThermoFisher Scientific Orbitrap Ascend. **a,** MS1 spectra of CALR and R-CALR using ITMS as a mass analyzer shows consistent charge state distribution as other instruments. **b,** Sequence coverage of unarginylated calreticulin when subjected to ETD or UVPD fragmentation. **c,** Annotated CALR and R-CALR MS2 spectra generated with ETD fragmentation. **d,** Annotated CALR and R-CALR deconvoluted MS2 spectra generated with UVPD fragmentation and deconvolved using FLASHDeconv.

| CALR and R-CALR Charge State 58 Mixing Experiment Signal | | | | |
|---|---|---|---|---|
| Sample | WT Peak Area (m/z 850.50) | R Peak Area (m/z 853.20) | % WT Signal | % R Signal |
| 02WT_08R | 177677 | 1333963 | 11.8 | 88.2 |
| 04WT_06R | 465622 | 916087 | 33.7 | 66.3 |
| 05WT_05R | 752091 | 769680 | 49.4 | 50.6 |
| 06WT_04R | 792489 | 559583 | 58.6 | 41.4 |
| 08WT_02R | 1130553 | 392557 | 74.2 | 25.8 |
| 10WT_01R | 460635 | 104218 | 81.5 | 18.5 |

**Supplemental Figure 5.** Peak area of *z* +58 CALR and R-CALR quantified over increasing ratios of WT and decreasing ratios of R. (**Related to Figure 4a**)

| 20 CALR Charges for XIC | | |
|---|---|---|
| **Charge** | **WT Predicted Average *m/z*** | **R Predicted Average *m/z*** |
| +45 | 1095.91 | 1099.38 |
| +46 | 1072.11 | 1075.51 |
| +47 | 1049.32 | 1052.65 |
| +48 | 1027.48 | 1030.74 |
| +49 | 1006.53 | 1009.72 |
| +50 | 986.42 | 989.55 |
| +51 | 967.10 | 970.16 |
| +52 | 948.52 | 951.53 |
| +53 | 930.65 | 933.59 |
| +54 | 913.43 | 916.32 |
| +55 | 896.84 | 899.68 |
| +56 | 880.84 | 883.63 |
| +57 | 865.41 | 868.15 |
| +58 | 850.50 | 853.20 |
| +59 | 836.11 | 838.75 |
| +60 | 822.19 | 824.79 |
| +61 | 808.73 | 811.29 |
| +62 | 795.70 | 798.22 |
| +63 | 783.08 | 785.56 |
| +64 | 770.86 | 773.30 |

**Supplemental Figure 6.** 20 charge states of CALR and R-CALR used in XIC for intact proteoform quantitation.

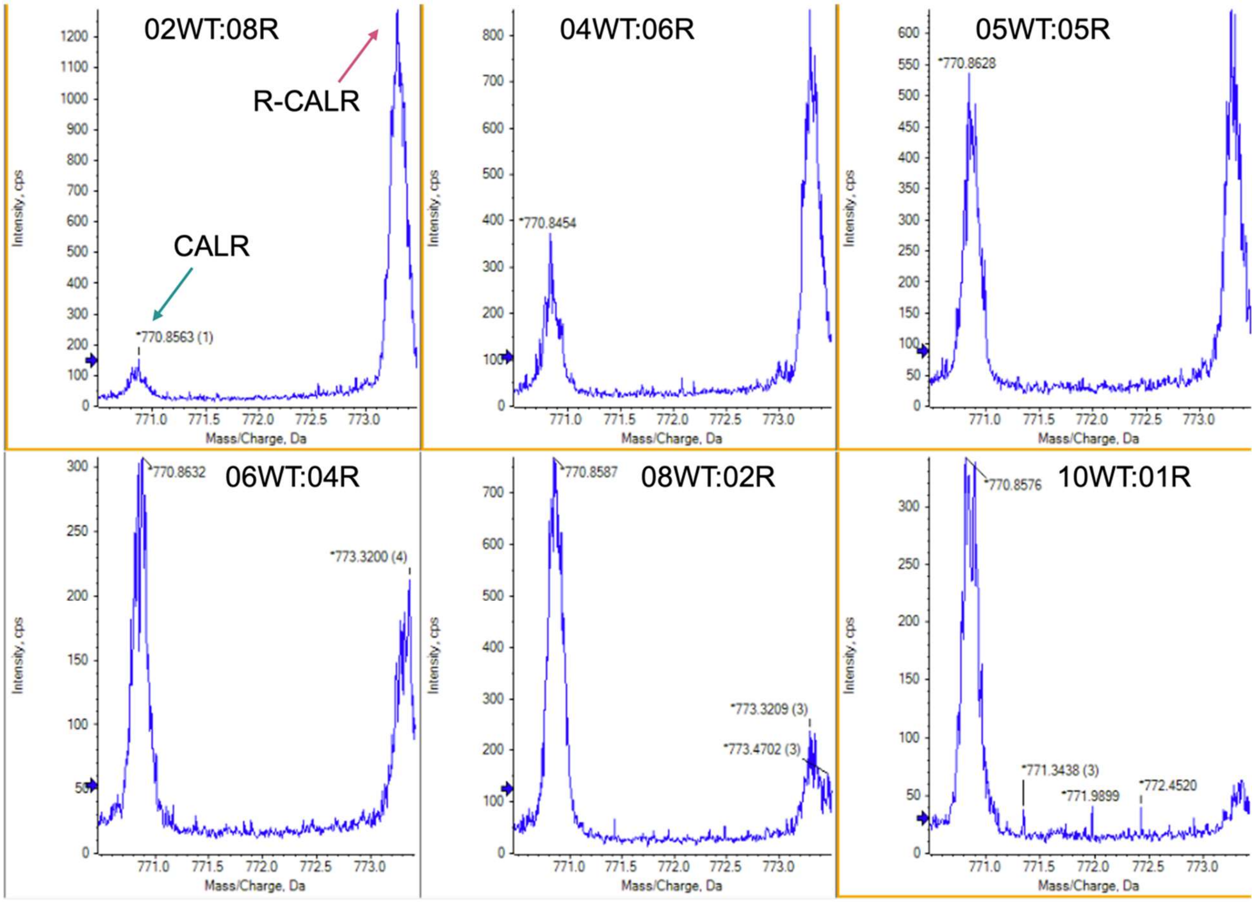


**Supplemental Figure 7.** Average spectra across the chromatographic peak of each CALR and R-CALR sample. At the MS1 spectral level, the shift in abundance is still observable.

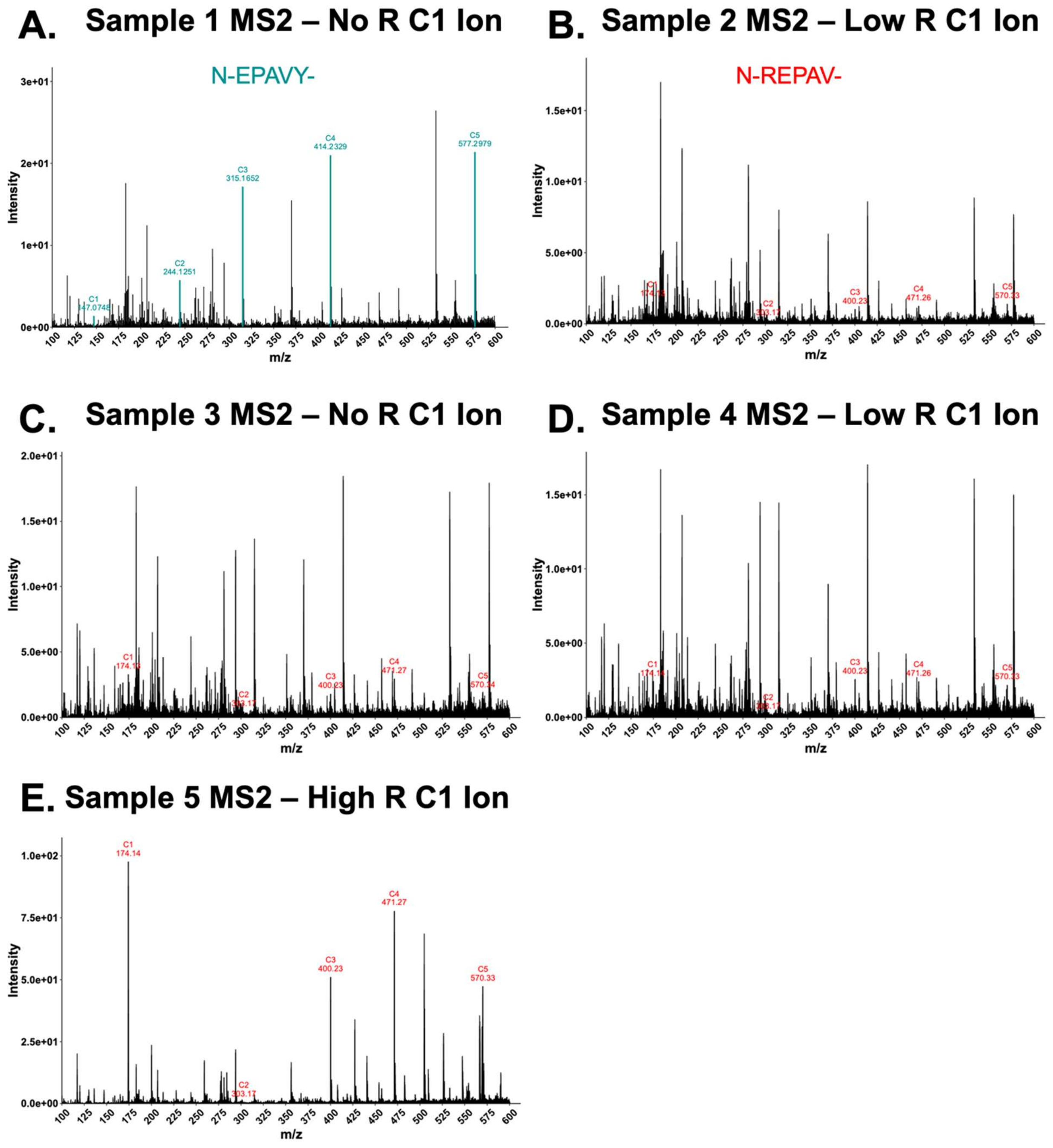


**Supplemental Figure 8.** EAD MS2 spectra of R-CALR targeting MS1 charge states at the noise level but still allow confident identification of R-CALR through the *c*-ion series. **a,** Sample 1 expressing calreticulin with no ATE1 co-expression shows the expected *c*-ion series, absent the signature *c1*-ion (*m/z* 174.13) of arginylated calreticulin. **b-d,** Samples 2 through 4 expressing calreticulin with ATE1 co-expression, showing low signal of the signature *c1*-ion (*m/z* 174.13) of arginylated calreticulin, confirming 18R presence. **e,** Sample 5 expressing calreticulin with extremely high ATE1 co-expression, showing a strong signal of the signature *c1*-ion (*m/z* 174.13) of arginylated calreticulin.

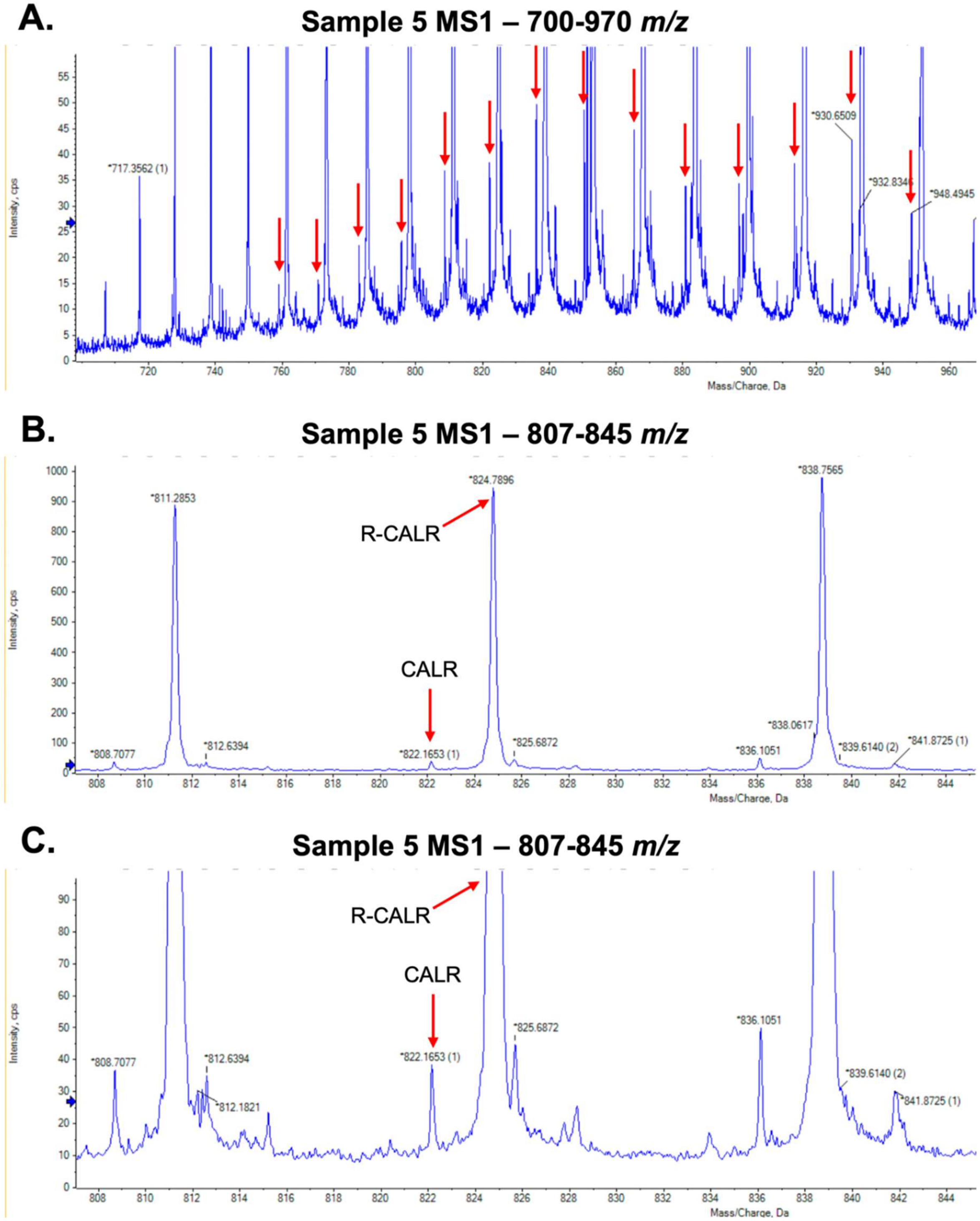


**Supplemental Figure 9.** MS1 spectra of Sample 5 R-CALR control sample, related to **Figure 1 and Figure 5**. **a,** MS1 spectra zoomed to *m/z* 700-970 and max intensity of 60 cps to enhance visualization of low-level CALR charge state distribution, indicated by red arrows. **b,** MS1 spectra zoomed to 807-845 *m/z* and max intensity of the spectra to enhance visualization of low-level CALR in relation to the high signal of R-CALR. **c,** MS1 spectra zoomed to 807-845 *m/z* and max intensity of 100 cps to enhance visualization of low-level CALR in relation to the high signal of R-CALR.

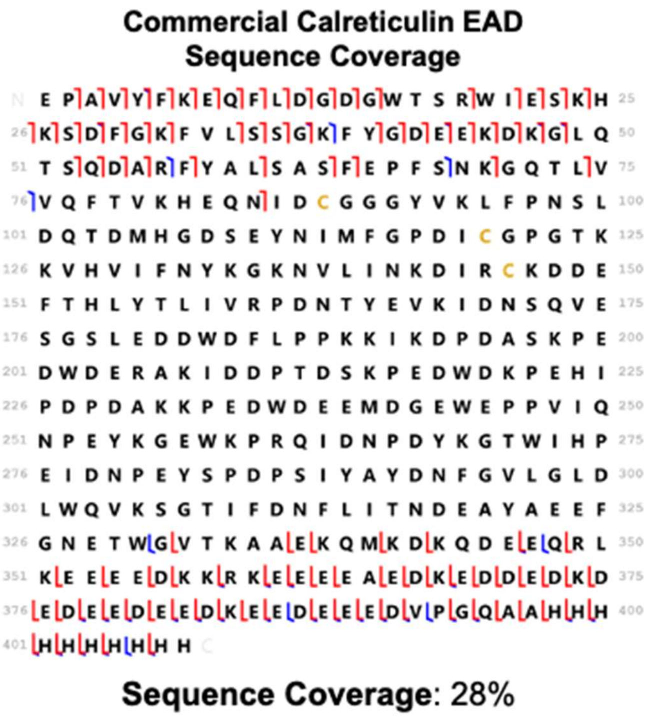


**Supplemental Figure 10.** EAD fragmentation sequence coverage map of commercial calreticulin with 8his *C*-terminal tag.

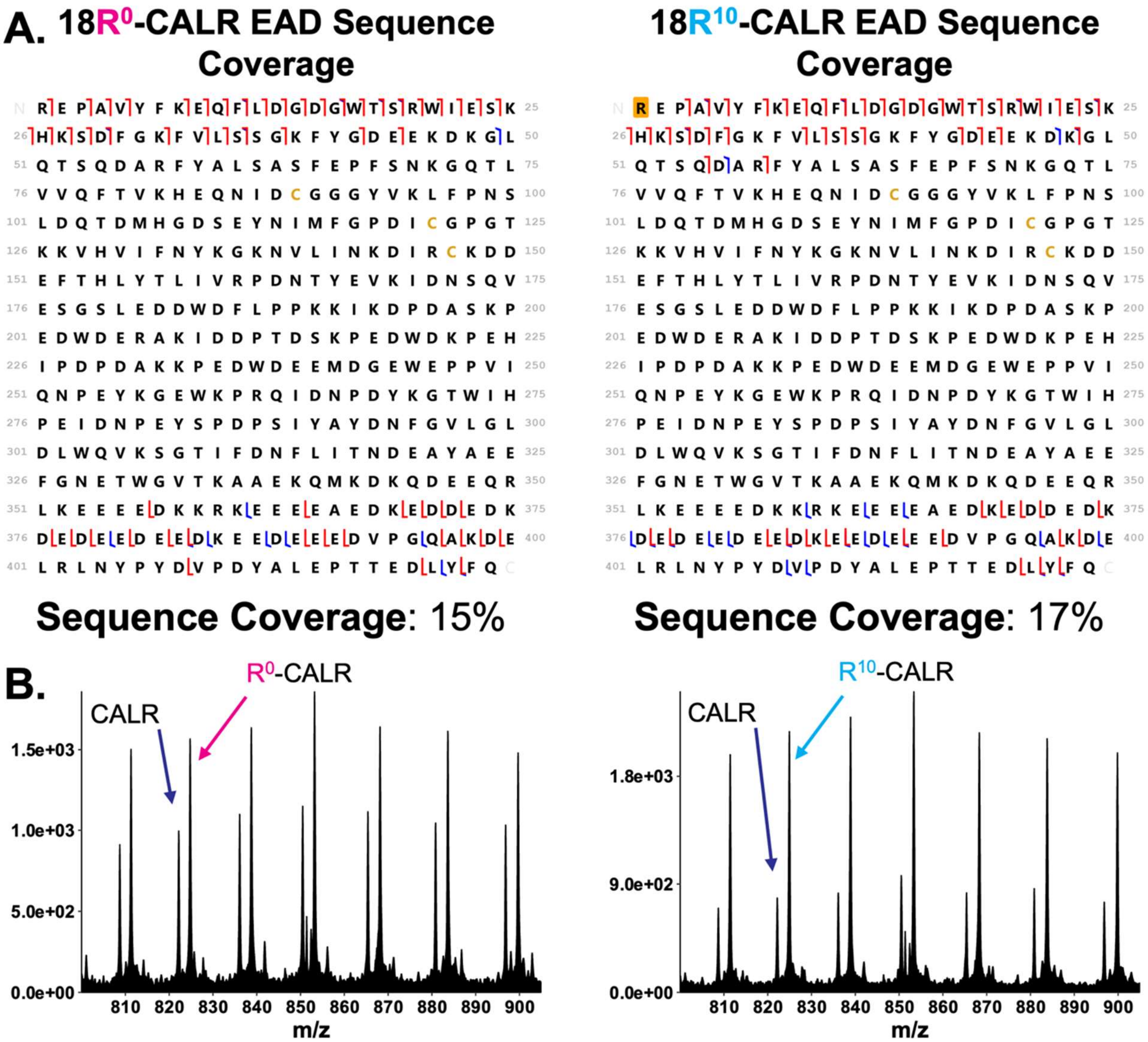


**Supplemental Figure 11. a,** Sequence coverage map of calreticulin labeled *N*-terminally with either 18R$^0$ or 18R$^{10}$ and affinity purified with Halo-tag then subjected to EAD fragmentation. The orange box around the *N*-terminal "R" in the 18R$^{10}$ calreticulin sequence coverage indicates a modification difference of 10.00824 Da was added to the arginine residue, corresponding to the mass shift of a heavy arginine from the light arginine. **b,** Focused view from *m/z* 800-900 of the sample containing CALR and 18R$^0$-CALR, or CALR and 18R$^{10}$-CALR. The unmodified CALR maintains consistent *m/z* measurement between samples, and a shift in *m/z* is observed between 18R$^0$ and 18R$^{10}$.

| CALR Charge States | | |
|---|---|---|
| Charge | Monoisotopic | Average |
| 55 | 896.30 | 896.84 |
| 56 | 880.31 | 880.84 |
| 57 | 864.89 | 865.41 |
| 58 | 849.99 | 850.50 |
| 59 | 835.60 | 836.11 |
| 60 | 821.69 | 822.19 |
| 61 | 808.24 | 808.73 |

| CALR R0 Charge States | | |
|---|---|---|
| Charge | Monoisotopic | Average |
| 55 | 899.14 | 899.68 |
| 56 | 883.10 | 883.63 |
| 57 | 867.62 | 868.15 |
| 58 | 852.68 | 853.20 |
| 59 | 838.25 | 838.75 |
| 60 | 824.29 | 824.79 |
| 61 | 810.80 | 811.29 |

| CALR R10 Charge States | | |
|---|---|---|
| Charge | Monoisotopic | Average |
| 55 | 899.32 | 899.86 |
| 56 | 883.28 | 883.81 |
| 57 | 867.80 | 868.32 |
| 58 | 852.86 | 853.37 |
| 59 | 838.42 | 838.92 |
| 60 | 824.46 | 824.96 |
| 61 | 810.96 | 811.45 |

| CALR *c* -ions | | |
|---|---|---|
| C+1 | C.Ion | AASeq |
| 147.0764 | 1 | E |
| 244.1292 | 2 | P |
| 315.1663 | 3 | A |
| 414.2347 | 4 | V |
| 577.2980 | 5 | Y |
| 724.3665 | 6 | F |
| 852.4614 | 7 | K |
| 981.5040 | 8 | E |
| 1109.5626 | 9 | Q |
| 1256.6310 | 10 | F |

| CALR R0 *c* -ions | | |
|---|---|---|
| C+1 | C.Ion | AASeq |
| 174.1349 | 1 | R |
| 303.1775 | 2 | E |
| 400.2303 | 3 | P |
| 471.2674 | 4 | A |
| 570.3358 | 5 | V |
| 733.3992 | 6 | Y |
| 880.4676 | 7 | F |
| 1008.5625 | 8 | K |
| 1137.6051 | 9 | E |
| 1265.6637 | 10 | Q |

| CALR R10 *c* -ions | | |
|---|---|---|
| C+1 | C.Ion | AASeq |
| 184.1432 | 1 | R |
| 313.1858 | 2 | E |
| 410.2385 | 3 | P |
| 481.2756 | 4 | A |
| 580.3441 | 5 | V |
| 743.4074 | 6 | Y |
| 890.4758 | 7 | F |
| 1018.5708 | 8 | K |
| 1147.6134 | 9 | E |
| 1275.6719 | 10 | Q |

**Supplemental Figure 12.** Predicted charge state distributions of WT-CALR, $R^0$-CALR, and $R^{10}$-CALR, and predicted *c*-ion fragment ion series following electron-based fragmentation.

**Plasmid sequence of CALR.**

gtatcttatcatgtctgtataccgtcgacctctagctagagcttggcgtaatcatggtcatagctgtttcctgtgtgaaattgttatccgctcacaattccaca
caacatacgagccggaagcataaagtgtaaagcctggggtgcctaatgagtgagctaactcacattaattgcgttgcgctcactgcccgctttccagt
cgggaaacctgtcgtgccagctgcattaatgaatcggccaacgcgcggggagaggcggtttgcgtattgggcgctcttccgcttcctcgctcactgac
tcgctgcgctcggtcgttcggctgcggcgagcggtatcagctcactcaaaggcggtaatacggttatccacagaatcaggggataacgcaggaaa
gaacatgtgagcaaaaggccagcaaaaggccaggaaccgtaaaaaggccgcgttgctggcgtttttccataggctccgcccccctgacgagcat
cacaaaaatcgacgctcaagtcagaggtggcgaaacccgacaggactataaagataccaggcgtttccccctggaagctccctcgtgcgctctcc
tgttccgaccctgccgcttaccggatacctgtccgcctttctcccttcgggaagcgtggcgctttctcatagctcacgctgtaggtatctcagttcggtgta
ggtcgttcgctccaagctgggctgtgtgcacgaaccccccgttcagcccgaccgctgcgccttatccggtaactatcgtcttgagtccaacccggtaa
gacacgacttatcgccactggcagcagccactggtaacaggattagcagagcgaggtatgtaggcggtgctacagagttcttgaagtggtggccta
actacggctacactagaaggacagtatttggtatctgcgctctgctgaagccagttaccttcggaaaaagagttggtagctcttgatccggcaaacaa
accaccgctggtagcggtggtttttttgtttgcaagcagcagattacgcgcagaaaaaaaggatctcaagaagatcctttgatcttttctacggggtctg
acgctcagtggaacgaaaactcacgttaagggattttggtcatgagattatcaaaaaggatcttcacctagatccttttaaattaaaaatgaagttttaa
atcaatctaaagtatatatgagtaaacttggtctgacagttaccaatgcttaatcagtgaggcacctatctcagcgatctgtctatttcgttcatccatagtt
gcctgactccccgtcgtgtagataactacgatacgggagggcttaccatctggccccagtgctgcaatgataccgcgagacccacgctcaccggct
ccagatttatcagcaataaaccagccagccggaagggccgagcgcagaagtggtcctgcaactttatccgcctccatccagtctattaattgttgccg
ggaagctagagtaagtagttcgccagttaatagtttgcgcaacgttgttgccattgctacaggcatcgtggtgtcacgctcgtcgtttggtatggcttcatt
cagctccggttcccaacgatcaaggcgagttacatgatcccccatgttgtgcaaaaaagcggttagctccttcggtcctccgatcgttgtcagaagta
agttggccgcagtgttatcactcatggttatggcagcactgcataattctcttactgtcatgccatccgtaagatgcttttctgtgactggtgagtactcaac
caagtcattctgagaatagtgtatgcggcgaccgagttgctcttgcccggcgtcaatacgggataataccgcgccacatagcagaactttaaaagtg
ctcatcattggaaaacgttcttcggggcgaaaactctcaaggatcttaccgctgttgagatccagttcgatgtaacccactcgtgcacccaactgatctt
cagcatcttttactttcaccagcgtttctgggtgagcaaaaacaggaaggcaaaatgccgcaaaaaagggaataagggcgacacggaaatgttg
aatactcatactcttcctttttcaatattattgaagcatttatcagggttattgtctcatgagcggatacatatttgaatgtatttagaaaaataaacaaatagg
ggttccgcgcacatttccccgaaaagtgccacctgacgtcgacggatcgggagatcttcaatattggccattagccatattattcattggttatatagcat
aaatcaatattggctattggccattgcatacgttgtatctatatcataatatgtacatttatattggctcatgtccaatatgaccgccatgttggcattgattatt
gactagttattaatagtaatcaattacggggtcattagttcatagcccatatatggagttccgcgttacataacttacggtaaatggcccgcctggctgac
cgcccaacgacccccgcccattgacgtcaataatgacgtatgttcccatagtaacgccaatagggactttccattgacgtcaatgggtggagtattta
cggtaaactgcccacttggcagtacatcaagtgtatcatatgccaagtccgccccctattgacgtcaatgacggtaaatggcccgcctggcattatgc
ccagtacatgaccttacgggactttcctacttggcagtacatctacgtattagtcatcgctattaccatggtgatgcggttttggcagtacaccaatgggc
gtggatagcggtttgactcacggggatttccaagtctccaccccattgacgtcaatgggagtttgttttggcaccaaaatcaacgggactttccaaaat
gtcgtaataaccccgccccgttgacgcaaatgggcggtaggcgtgtacggtgggaggtctatataagcagagctggtttagtgaaccgtcagatca
ctagaagctttattgcggtagtttatcacagttaaattgctaacgcagtcagtgcttctgacacaacagtctcgaacttaagctgcagaagttggtcgtga
ggcactgggcaggtaagtatcaaggttacaagacaggtttaaggagaccaatagaaactgggcttgtcgagacagagaagactcttgcgtttctga
taggcacctattggtcttactgacatccactttgcctttctctccacaggtgtccactcccagttcaattacagctcttaaggctagagtattaatacgactc
actatagggctagcgatcgcgccaccATGCTGCTATCCGTGCCGCTGCTGCTCGGCCTCCTCGGCCTGGCCGTC
GCCGAGCCTGCCGTCTACTTCAAGGAGCAGTTTCTGGACGGAGACGGGTGGACTTCCCGCTGGATC
GAATCCAAACACAAGTCAGATTTTGGCAAATTCGTTCTCAGTTCCGGCAAGTTCTACGGTGACGAGGA
GAAAGATAAAGGTTTGCAGACAAGCCAGGATGCACGCTTTTATGCTCTGTCGGCCAGTTTCGAGCCTT
TCAGCAACAAAGGCCAGACGCTGGTGGTGCAGTTCACGGTGAAACATGAGCAGAACATCGACTGTG
GGGGCGGCTATGTGAAGCTGTTTCCTAATAGTTTGGACCAGACAGACATGCACGGAGACTCAGAATA
CAACATCATGTTTGGTCCCGACATCTGTGGCCCTGGCACCAAGAAGGTTCATGTCATCTTCAACTACA
AGGGCAAGAACGTGCTGATCAACAAGGACATCCGTTGCAAGGATGATGAGTTTACACACCTGTACAC
ACTGATTGTGCGGCCAGACAACACCTATGAGGTGAAGATTGACAACAGCCAGGTGGAGTCCGGCTC
CTTGGAAGACGATTGGGACTTCCTGCCACCCAAGAAGATAAAGGATCCTGATGCTTCAAAACCGGAA
GACTGGGATGAGCGGGCCAAGATCGATGATCCCACAGACTCCAAGCCTGAGGACTGGGACAAGCCC
GAGCATATCCCTGACCCTGATGCTAAGAAGCCCGAGGACTGGGATGAAGAGATGGACGGAGAGTGG
GAACCCCCAGTGATTCAGAACCCTGAGTACAAGGGTGAGTGGAAGCCCCGGCAGATCGACAACCCA
GATTACAAGGGCACTTGGATCCACCCAGAAATTGACAACCCCGAGTATTCTCCCGATCCCAGTATCTAT
GCCTATGATAACTTTGGCGTGCTGGGCCTGGACCTCTGGCAGGTCAAGTCTGGCACCATCTTTGACA
ACTTCCTCATCACCAACGATGAGGCATACGCTGAGGAGTTTGGCAACGAGACGTGGGGCGTAACAAA
GGCAGCAGAGAAACAAATGAAGGACAAACAGGACGAGGAGCAGAGGCTTaaggaggaggaagaagacaag
aaacgcaaagaggaggaggaggcagaggacaaggaggatgatgaggacaaagatgaggatgaggaggatgaggaggacaaggaggaa
gatgaggaggaagatgTCCCCGGCCAGGCCAAGGACGAGCTGcgtttaaactacccatacgatgttccagattacgctctcgag
ccaaccactgaggatctgtactttcagagcgataacgatggatccgaaatcggtactggctttccattcgacccccattatgtggaagtcctgggcga
gcgcatgcactacgtcgatgttggtccgcgcgatggcacccctgtgctgttcctgcacggtaacccgacctcctcctacgtgtggcgcaacatcatcc
cgcatgttgcaccgacccatcgctgcattgctccagacctgatcggtatgggcaaatccgacaaaccagacctgggttatttcttcgacgaccacgtc

cgcttcatggatgccttcatcgaagccctgggtctggaagaggtcgtcctggtcattcacgactggggctccgctctgggtttccactgggccaagcgc
aatccagagcgcgtcaaaggtattgcatttatggagttcatccgccctatcccgacctgggacgaatggccagaatttgcccgcgagaccttccagg
ccttccgcaccaccgacgtcggccgcaagctgatcatcgatcagaacgtttttatcgagggtacgctgccgatgggtgtcgtccgcccgctgactga
agtcgagatggaccattaccgcgagccgttcctgaatcctgttgaccgcgagccactgtggcgcttcccaaacgagctgccaatcgccggtgagcc
agcgaacatcgtcgcgctggtcgaagaatacatggactggctgcaccagtcccctgtcccgaagctgctgttctggggcaccccaggcgttctgatc
ccaccggccgaagccgctcgcctggccaaaagcctgcctaactgcaaggctgtggacatcggcccgggtctgaatctgctgcaagaagacaac
ccggacctgatcggcagcgagatcgcgcgctggctgtctactctggagatttccggttaatagaattctagagtcgacctgcaggcatgcaagctgat
ccggctgctaacaaagcccgaaaggaagctgagttggctgctgccaccgctgagcaataactagcataaccccttggggcggccaaacccgctg
atcagcctcgactgtgccttctagttgccagccatctgttgtttgcccctcccccgtgccttccttgaccctggaaggtgccactcccactgtcctttcctaat
aaaatgaggaaattgcatcgcattgtctgagtaggtgtcattctattctggggggtggggtggggcaggacagcaagggggaggattgggaagac
aatagcaggcatgctggggatgcggtgggctctatggcttctgaggcggaaagaaccagctggggctctagggggtatccccacgcgccctgtag
cggcgcattaagcgcggcgggtgtggtggttacgcgcagcgtgaccgctacacttgccagcgccctagcgcccgctcctttcgctttcttcccttccttt
ctcgccacgttcgccggctttccccgtcaagctctaaatcgggggctccctttagggttccgatttagtgctttacggcacctcgaccccaaaaaacttg
attagggtgatggttcacgtacctagaagttcctattccgaagttcctattctctagaaagtataggaacttccttggccaaaaagcctgaactcaccgc
gacgtctgtcgagaagtttctgatcgaaaagttcgacagcgtctccgacctgatgcagctctcggagggcgaagaatctcgtgctttcagcttcgatgt
aggagggcgtggatatgtcctgcgggtaaatagctgcgccgatggtttctacaaagatcgttatgtttatcggcactttgcatcggccgcgctcccgatt
ccggaagtgcttgacattggggaattcagcgagagcctgacctattgcatctcccgccgtgcacagggtgtcacgttgcaagacctgcctgaaacc
gaactgcccgctgttctgcagccggtcgcggaggccatggatgctatcgctgcggccgatcttagccagacgagcgggttcggcccattcggaccg
caaggaatcggtcaatacactacatggcgtgatttcatatgcgcgattgctgatccccatgtgtatcactggcaaactgtgatggacgacaccgtcagt
gcgtccgtcgcgcaggctctcgatgagctgatgctttgggccgaggactgccccgaagtccggcacctcgtgcacgcggatttcggctccaacaat
gtcctgacggacaatggccgcataacagcggtcattgactggagcgaggcgatgttcggggattcccaatacgaggtcgccaacatcttcttctgga
ggccgtggttggcttgtatggagcagcagacgcgctacttcgagcggaggcatccggagcttgcaggatcgccgcggctccgggcgtatatgctcc
gcattggtcttgaccaactctatcagagcttggttgacggcaatttcgatgatgcagcttgggcgcagggtcgatgcgacgcaatcgtccgatccgga
gccgggactgtcgggcgtacacaaatcgcccgcagaagcgcggccgtctggaccgatggctgtgtagaagtactcgccgatagtggaaaccga
cgccccagcactcgtccgagggcaaaggaatagcacgtactacgagatttcgattccaccgccgccttctatgaaaggttgggcttcggaatcgtttt
ccgggacgccggctggatgatcctccagcgcggggatctcatgctggagttcttcgcccaccccaacttgtttattgcagcttataatggttacaaata
aagcaatagcatcacaaatttcacaaataaagcatttttttcactgcattctagttgtggtttgtccaaactcatcaat

**Plasmid sequence of R-CALR.**

…ATGCTGCTATCCGTGCCGCTGCTGCTCGGCCTCCTCGGCCTGGCCGTCGCCAGAGAGCCTGCC…

**Plasmid sequence of ATE1_FLAG.**

GACGGATCGGGAGATCTCCCGATCCCCTATGGTGCACTCTCAGTACAATCTGCTCTGATGCCGCATAG
TTAAGCCAGTATCTGCTCCCTGCTTGTGTGTTGGAGGTCGCTGAGTAGTGCGCGAGCAAAATTTAAGC
TACAACAAGGCAAGGCTTGACCGACAATTGCATGAAGAATCTGCTTAGGGTTAGGCGTTTTGCGCTG
CTTCGCGATGTACGGGCCAGATATACGCGTTGACATTGATTATTGACTAGTTATTAATAGTAATCAATTAC
GGGGTCATTAGTTCATAGCCCATATATGGAGTTCCGCGTTACATAACTTACGGTAAATGGCCCGCCTGG
CTGACCGCCCAACGACCCCCGCCCATTGACGTCAATAATGACGTATGTTCCCATAGTAACGCCAATAG
GGACTTTCCATTGACGTCAATGGGTGGAGTATTTACGGTAAACTGCCCACTTGGCAGTACATCAAGTG
TATCATATGCCAAGTACGCCCCCTATTGACGTCAATGACGGTAAATGGCCCGCCTGGCATTATGCCCA
GTACATGACCTTATGGGACTTTCCTACTTGGCAGTACATCTACGTATTAGTCATCGCTATTACCATGGTG
ATGCGGTTTTGGCAGTACATCAATGGGCGTGGATAGCGGTTTGACTCACGGGGATTTCCAAGTCTCC
ACCCCATTGACGTCAATGGGAGTTTGTTTTGGCACCAAAATCAACGGGACTTTCCAAAATGTCGTAAC
AACTCCGCCCCATTGACGCAAATGGGCGGTAGGCGTGTACGGTGGGAGGTCTATATAAGCAGAGCTC
TCTGGCTAACTAGAGAACCCACTGCTTACTGGCTTATCGAAATTAATACGACTCACTATAGGGAGACCC
AAGCTGGCTAGCGTTTAAACTTAAGCTTGGTACCGAGCTCGGATCCGCCACCATGGCTTTCTGGGCG
GGGGGTTCGCCCAGCGTCGTGGACTATTTCCCTAGCGAGGACTTCTACCGCTGCGGCTACTGCAAG
AACGAGTCGGGCAGCCGCTCCAATGGCATGTGGGCACATTCCATGACAGTACAGGATTATCAGGATC
TCATAGACCGAGGATGGCGAAGAAGTGGAAAATATGTGTACAAACCTGTCATGAATCAAACATGTTGT
CCTCAGTACACAATAAGGTGCCGACCTTTACAATTTCAGCCTTCAAAATCTCACAAGAAGGTTTTGAAA
AAAATGTTGAAATTTCTAGCTAAAGGGGAGGTTCCCAAAGGAAGTTGTGAGGATGAGCCCATGGATTC
CACAATGGATGATGCTGTTGCGGGTGACTTTGCATTGATAAATAAACTGGATATACAGTGTGATCTTAAA
ACACTCAGTGATGACATCAAAGAGAGTTTAGAGAGTGAAGGAAAAAATTCAAAGAAAGAAGAACCTCA
GGAATTACTTCAGTCACAAGATTTCGTAGGAGAGAAGTTGGGCTCTGGTGAACCGTCACATTCAGTTA
AAGTTCACACAGTTCCTAAGCCAGGCAAAGGGGCTGATTTGAGTAAGCCTCCATGTCGAAAAGCAAA
GGAAATCCGGAAAGAAAGGAAAAGGTTAAAACTAATGCAGCAGAACCCAGCTGGAGAACTTGAGGGT
TTCCAGGCTCAAGGTCACCCACCATCTTTGTTTCCACCAAAGGCTAAATCCAACCAGCCAAAATCACT
CGAAGATTTAATTTTTGAGTCTTTACCAGAGAATGCATCACACAAGTTAGAGGTGAGGGTGGTGAGAT

CATCTCCACCAAGTTCGCAGTTCAAAGCCACACTTCTGGAGTCTTACCAGGTCTATAAACGTTACCAG
ATGGTTATTCACAAGAACCCACCTGATACGCCAACCGAAAGCCAGTTCACAAGATTCCTTTGCAGTTC
ACCCTTGGAGGCAGAGACTCCCCCTAATGGGCCAGATTGTGGCTATGGCTCCTTTCACCAGCAGTAC
TGGCTTGACGGAAAGATCATTGCTGTGGGGGTGATTGACATCCTCCCAAACTGTGTATCATCTGTGTA
TTTGTACTACGATCCTGATTATTCGTTTTTGTCTTTGGGCGTCTACTCTGCACTACGAGAAATTGCTTTT
ACTAGGCAGCTTCATGAGAAAACTTCTCAACTCAGCTATTATTATATGGGTTTCTACATTCATTCATGTC
CCAAGATGAAATATAAGGGTCAGTATAGACCTTCTGATTTGCTGTGCCCTGAGACATATGTTTGGGTAC
CCATTGAGCAATGCCTGCCTTCACTTGAAAACTCCAAGTACTGCCGTTTCAACCAGGACCCAGAAGC
AGTGGATGAGGATCGCAGTACGGAACCTGACCGATTGCAGGTGTTTCACAAGAGAGCCATCATGCCT
TACGGTGTTTATAAGAAACAGCAGAAAGACCCAAGTGAGGAGGCTGCTGTTCTGCAGTACGCCAGCC
TGGTGGGGCAGAAGTGCTCCGAGCGGATGCTGCTGTTCAGAAACGATTACAAGGATGACGACGATAA
GTGATAAACCCGCTGATCAGCCTCGACTGTGCCTTCTAGTTGCCAGCCATCTGTTGTTTGCCCCTCCC
CCGTGCCTTCCTTGACCCTGGAAGGTGCCACTCCCACTGTCCTTTCCTAATAAAATGAGGAAATTGCA
TCGCATTGTCTGAGTAGGTGTCATTCTATTCTGGGGGGTGGGGTGGGGCAGGACAGCAAGGGGGAG
GATTGGGAAGACAATAGCAGGCATGCTGGGGATGCGGTGGGCTCTATGGCTTCTGAGGCGGAAAGA
ACCAGCTGGGGCTCTAGGGGGTATCCCCACGCGCCCTGTAGCGGCGCATTAAGCGCGGCGGGTGT
GGTGGTTACGCGCAGCGTGACCGCTACACTTGCCAGCGCCCTAGCGCCCGCTCCTTTCGCTTTCTT
CCCTTCCTTTCTCGCCACGTTCGCCGGCTTTCCCCGTCAAGCTCTAAATCGGGGGCTCCCTTTAGGG
TTCCGATTTAGTGCTTTACGGCACCTCGACCCCAAAAAACTTGATTAGGGTGATGGTTCACGTAGTGG
GCCATCGCCCTGATAGACGGTTTTTCGCCCTTTGACGTTGGAGTCCACGTTCTTTAATAGTGGACTCT
TGTTCCAAACTGGAACAACACTCAACCCTATCTCGGTCTATTCTTTTGATTTATAAGGGATTTTGCCGAT
TTCGGCCTATTGGTTAAAAAATGAGCTGATTTAACAAAAATTTAACGCGAATTAATTCTGTGGAATGTGT
GTCAGTTAGGGTGTGGAAAGTCCCCAGGCTCCCCAGCAGGCAGAAGTATGCAAAGCATGCATCTCAA
TTAGTCAGCAACCAGGTGTGGAAAGTCCCCAGGCTCCCCAGCAGGCAGAAGTATGCAAAGCATGCAT
CTCAATTAGTCAGCAACCATAGTCCCGCCCCTAACTCCGCCCATCCCGCCCCTAACTCCGCCCAGTT
CCGCCCATTCTCCGCCCCATGGCTGACTAATTTTTTTTATTTATGCAGAGGCCGAGGCCGCCTCTGCC
TCTGAGCTATTCCAGAAGTAGTGAGGAGGCTTTTTTGGAGGCCTAGGCTTTTGCAAAAAGCTCCCGG
GAGCTTGTATATCCATTTTCGGATCTGATCAAGAGACAGGATGAGGATCGTTTCGCATGATTGAACAAG
ATGGATTGCACGCAGGTTCTCCGGCCGCTTGGGTGGAGAGGCTATTCGGCTATGACTGGGCACAAC
AGACAATCGGCTGCTCTGATGCCGCCGTGTTCCGGCTGTCAGCGCAGGGGCGCCCGGTTCTTTTTG
TCAAGACCGACCTGTCCGGTGCCCTGAATGAACTGCAGGACGAGGCAGCGCGGCTATCGTGGCTGG
CCACGACGGGCGTTCCTTGCGCAGCTGTGCTCGACGTTGTCACTGAAGCGGGAAGGGACTGGCTG
CTATTGGGCGAAGTGCCGGGGCAGGATCTCCTGTCATCTCACCTTGCTCCTGCCGAGAAAGTATCCA
TCATGGCTGATGCAATGCGGCGGCTGCATACGCTTGATCCGGCTACCTGCCCATTCGACCACCAAGC
GAAACATCGCATCGAGCGAGCACGTACTCGGATGGAAGCCGGTCTTGTCGATCAGGATGATCTGGAC
GAAGAGCATCAGGGGCTCGCGCCAGCCGAACTGTTCGCCAGGCTCAAGGCGCGCATGCCCGACGG
CGAGGATCTCGTCGTGACCCATGGCGATGCCTGCTTGCCGAATATCATGGTGGAAAATGGCCGCTTT
TCTGGATTCATCGACTGTGGCCGGCTGGGTGTGGCGGACCGCTATCAGGACATAGCGTTGGCTACC
CGTGATATTGCTGAAGAGCTTGGCGGCGAATGGGCTGACCGCTTCCTCGTGCTTTACGGTATCGCCG
CTCCCGATTCGCAGCGCATCGCCTTCTATCGCCTTCTTGACGAGTTCTTCTGAGCGGGACTCTGGGG
TTCGAAATGACCGACCAAGCGACGCCCAACCTGCCATCACGAGATTTCGATTCCACCGCCGCCTTCT
ATGAAAGGTTGGGCTTCGGAATCGTTTTCCGGGACGCCGGCTGGATGATCCTCCAGCGCGGGGATC
TCATGCTGGAGTTCTTCGCCCACCCCAACTTGTTTATTGCAGCTTATAATGGTTACAAATAAAGCAATAG
CATCACAAATTTCACAAATAAAGCATTTTTTTCACTGCATTCTAGTTGTGGTTTGTCCAAACTCATCAAT
GTATCTTATCATGTCTGTATACCGTCGACCTCTAGCTAGAGCTTGGCGTAATCATGGTCATAGCTGTTTC
CTGTGTGAAATTGTTATCCGCTCACAATTCCACACAACATACGAGCCGGAAGCATAAAGTGTAAAGCC
TGGGGTGCCTAATGAGTGAGCTAACTCACATTAATTGCGTTGCGCTCACTGCCCGCTTTCCAGTCGG
GAAACCTGTCGTGCCAGCTGCATTAATGAATCGGCCAACGCGCGGGGAGAGGCGGTTTGCGTATTG
GGCGCTCTTCCGCTTCCTCGCTCACTGACTCGCTGCGCTCGGTCGTTCGGCTGCGGCGAGCGGTAT
CAGCTCACTCAAAGGCGGTAATACGGTTATCCACAGAATCAGGGGATAACGCAGGAAAGAACATGTG
AGCAAAAGGCCAGCAAAAGGCCAGGAACCGTAAAAAGGCCGCGTTGCTGGCGTTTTTCCATAGGCT
CCGCCCCCCTGACGAGCATCACAAAAATCGACGCTCAAGTCAGAGGTGGCGAAACCCGACAGGACT
ATAAAGATACCAGGCGTTTCCCCCTGGAAGCTCCCTCGTGCGCTCTCCTGTTCCGACCCTGCCGCTT
ACCGGATACCTGTCCGCCTTTCTCCCTTCGGGAAGCGTGGCGCTTTCTCATAGCTCACGCTGTAGGT
ATCTCAGTTCGGTGTAGGTCGTTCGCTCCAAGCTGGGCTGTGTGCACGAACCCCCCGTTCAGCCCG
ACCGCTGCGCCTTATCCGGTAACTATCGTCTTGAGTCCAACCCGGTAAGACACGACTTATCGCCACTG

GCAGCAGCCACTGGTAACAGGATTAGCAGAGCGAGGTATGTAGGCGGTGCTACAGAGTTCTTGAAGT
GGTGGCCTAACTACGGCTACACTAGAAGAACAGTATTTGGTATCTGCGCTCTGCTGAAGCCAGTTACC
TTCGGAAAAAGAGTTGGTAGCTCTTGATCCGGCAAACAAACCACCGCTGGTAGCGGTGGTTTTTTTG
TTTGCAAGCAGCAGATTACGCGCAGAAAAAAAGGATCTCAAGAAGATCCTTTGATCTTTTCTACGGGG
TCTGACGCTCAGTGGAACGAAAACTCACGTTAAGGGATTTTGGTCATGAGATTATCAAAAAGGATCTT
CACCTAGATCCTTTTAAATTAAAAATGAAGTTTTAAATCAATCTAAAGTATATATGAGTAAACTTGGTCTG
ACAGTTACCAATGCTTAATCAGTGAGGCACCTATCTCAGCGATCTGTCTATTTCGTTCATCCATAGTTG
CCTGACTCCCCGTCGTGTAGATAACTACGATACGGGAGGGCTTACCATCTGGCCCCAGTGCTGCAAT
GATACCGCGAGACCCACGCTCACCGGCTCCAGATTTATCAGCAATAAACCAGCCAGCCGGAAGGGC
CGAGCGCAGAAGTGGTCCTGCAACTTTATCCGCCTCCATCCAGTCTATTAATTGTTGCCGGGAAGCTA
GAGTAAGTAGTTCGCCAGTTAATAGTTTGCGCAACGTTGTTGCCATTGCTACAGGCATCGTGGTGTCA
CGCTCGTCGTTTGGTATGGCTTCATTCAGCTCCGGTTCCCAACGATCAAGGCGAGTTACATGATCCCC
CATGTTGTGCAAAAAAGCGGTTAGCTCCTTCGGTCCTCCGATCGTTGTCAGAAGTAAGTTGGCCGCA
GTGTTATCACTCATGGTTATGGCAGCACTGCATAATTCTCTTACTGTCATGCCATCCGTAAGATGCTTTT
CTGTGACTGGTGAGTACTCAACCAAGTCATTCTGAGAATAGTGTATGCGGCGACCGAGTTGCTCTTGC
CCGGCGTCAATACGGGATAATACCGCGCCACATAGCAGAACTTTAAAAGTGCTCATCATTGGAAAACG
TTCTTCGGGGCGAAAACTCTCAAGGATCTTACCGCTGTTGAGATCCAGTTCGATGTAACCCACTCGTG
CACCCAACTGATCTTCAGCATCTTTTACTTTCACCAGCGTTTCTGGGTGAGCAAAAACAGGAAGGCAA
AATGCCGCAAAAAAGGGAATAAGGGCGACACGGAAATGTTGAATACTCATACTCTTCCTTTTTCAATAT
TATTGAAGCATTTATCAGGGTTATTGTCTCATGAGCGGATACATATTTGAATGTATTTAGAAAAATAAACA
AATAGGGGTTCCGCGCACATTTCCCCGAAAAGTGCCACCTGACGTC
**Plasmid sequence of empty pcDNA3.1+/C-(k)DYK plasmid.**
GACGGATCGGGAGATCTCCCGATCCCCTATGGTGCACTCTCAGTACAATCTGCTCTGATGCCGCATAG
TTAAGCCAGTATCTGCTCCCTGCTTGTGTGTTGGAGGTCGCTGAGTAGTGCGCGAGCAAAATTTAAGC
TACAACAAGGCAAGGCTTGACCGACAATTGCATGAAGAATCTGCTTAGGGTTAGGCGTTTTGCGCTG
CTTCGCGATGTACGGGCCAGATATACGCGTTGACATTGATTATTGACTAGTTATTAATAGTAATCAATTAC
GGGGTCATTAGTTCATAGCCCATATATGGAGTTCCGCGTTACATAACTTACGGTAAATGGCCCGCCTGG
CTGACCGCCCAACGACCCCCGCCCATTGACGTCAATAATGACGTATGTTCCCATAGTAACGCCAATAG
GGACTTTCCATTGACGTCAATGGGTGGAGTATTTACGGTAAACTGCCCACTTGGCAGTACATCAAGTG
TATCATATGCCAAGTACGCCCCCTATTGACGTCAATGACGGTAAATGGCCCGCCTGGCATTATGCCCA
GTACATGACCTTATGGGACTTTCCTACTTGGCAGTACATCTACGTATTAGTCATCGCTATTACCATGGTG
ATGCGGTTTTGGCAGTACATCAATGGGCGTGGATAGCGGTTTGACTCACGGGGATTTCCAAGTCTCC
ACCCCATTGACGTCAATGGGAGTTTGTTTTGGCACCAAAATCAACGGGACTTTCCAAAATGTCGTAAC
AACTCCGCCCCATTGACGCAAATGGGCGGTAGGCGTGTACGGTGGGAGGTCTATATAAGCAGAGCTC
TCTGGCTAACTAGAGAACCCACTGCTTACTGGCTTATCGAAATTAATACGACTCACTATAGGGAGACCC
AAGCTGGCTAGCGTTTAAACTTAAGCTTGGTACCGAGCTCGGATCCGCCACCGAATTCTGCAGATATC
CAGCACAGTGGCGGCCGCTCGAGTCTAGAGGGCCCGATTACAAGGATGACGACGATAAGTGATAAAC
CCGCTGATCAGCCTCGACTGTGCCTTCTAGTTGCCAGCCATCTGTTGTTTGCCCCTCCCCCGTGCCT
TCCTTGACCCTGGAAGGTGCCACTCCCACTGTCCTTTCCTAATAAAATGAGGAAATTGCATCGCATTG
TCTGAGTAGGTGTCATTCTATTCTGGGGGGTGGGGTGGGGCAGGACAGCAAGGGGGAGGATTGGGA
AGACAATAGCAGGCATGCTGGGGATGCGGTGGGCTCTATGGCTTCTGAGGCGGAAAGAACCAGCTG
GGGCTCTAGGGGGTATCCCCACGCGCCCTGTAGCGGCGCATTAAGCGCGGCGGGTGTGGTGGTTAC
GCGCAGCGTGACCGCTACACTTGCCAGCGCCCTAGCGCCCGCTCCTTTCGCTTTCTTCCCTTCCTTT
CTCGCCACGTTCGCCGGCTTTCCCCGTCAAGCTCTAAATCGGGGGCTCCCTTTAGGGTTCCGATTTA
GTGCTTTACGGCACCTCGACCCCAAAAAACTTGATTAGGGTGATGGTTCACGTAGTGGGCCATCGCC
CTGATAGACGGTTTTTCGCCCTTTGACGTTGGAGTCCACGTTCTTTAATAGTGGACTCTTGTTCCAAAC
TGGAACAACACTCAACCCTATCTCGGTCTATTCTTTTGATTTATAAGGGATTTTGCCGATTTCGGCCTAT
TGGTTAAAAAATGAGCTGATTTAACAAAAATTTAACGCGAATTAATTCTGTGGAATGTGTGTCAGTTAGG
GTGTGGAAAGTCCCCAGGCTCCCCAGCAGGCAGAAGTATGCAAAGCATGCATCTCAATTAGTCAGCA
ACCAGGTGTGGAAAGTCCCCAGGCTCCCCAGCAGGCAGAAGTATGCAAAGCATGCATCTCAATTAGT
CAGCAACCATAGTCCCGCCCCTAACTCCGCCCATCCCGCCCCTAACTCCGCCCAGTTCCGCCCATTC
TCCGCCCCATGGCTGACTAATTTTTTTTATTTATGCAGAGGCCGAGGCCGCCTCTGCCTCTGAGCTAT
TCCAGAAGTAGTGAGGAGGCTTTTTTGGAGGCCTAGGCTTTTGCAAAAAGCTCCCGGGAGCTTGTAT
ATCCATTTTCGGATCTGATCAAGAGACAGGATGAGGATCGTTTCGCATGATTGAACAAGATGGATTGC
ACGCAGGTTCTCCGGCCGCTTGGGTGGAGAGGCTATTCGGCTATGACTGGGCACAACAGACAATCG
GCTGCTCTGATGCCGCCGTGTTCCGGCTGTCAGCGCAGGGGCGCCCGGTTCTTTTTGTCAAGACCG

ACCTGTCCGGTGCCCTGAATGAACTGCAGGACGAGGCAGCGCGGCTATCGTGGCTGGCCACGACG
GGCGTTCCTTGCGCAGCTGTGCTCGACGTTGTCACTGAAGCGGGAAGGGACTGGCTGCTATTGGGC
GAAGTGCCGGGGCAGGATCTCCTGTCATCTCACCTTGCTCCTGCCGAGAAAGTATCCATCATGGCTG
ATGCAATGCGGCGGCTGCATACGCTTGATCCGGCTACCTGCCCATTCGACCACCAAGCGAAACATCG
CATCGAGCGAGCACGTACTCGGATGGAAGCCGGTCTTGTCGATCAGGATGATCTGGACGAAGAGCAT
CAGGGGCTCGCGCCAGCCGAACTGTTCGCCAGGCTCAAGGCGCGCATGCCCGACGGCGAGGATCT
CGTCGTGACCCATGGCGATGCCTGCTTGCCGAATATCATGGTGGAAAATGGCCGCTTTTCTGGATTCA
TCGACTGTGGCCGGCTGGGTGTGGCGGACCGCTATCAGGACATAGCGTTGGCTACCCGTGATATTGC
TGAAGAGCTTGGCGGCGAATGGGCTGACCGCTTCCTCGTGCTTTACGGTATCGCCGCTCCCGATTCG
CAGCGCATCGCCTTCTATCGCCTTCTTGACGAGTTCTTCTGAGCGGGACTCTGGGGTTCGAAATGAC
CGACCAAGCGACGCCCAACCTGCCATCACGAGATTTCGATTCCACCGCCGCCTTCTATGAAAGGTTG
GGCTTCGGAATCGTTTTCCGGGACGCCGGCTGGATGATCCTCCAGCGCGGGGATCTCATGCTGGAG
TTCTTCGCCCACCCCAACTTGTTTATTGCAGCTTATAATGGTTACAAATAAAGCAATAGCATCACAAATT
TCACAAATAAAGCATTTTTTTCACTGCATTCTAGTTGTGGTTTGTCCAAACTCATCAATGTATCTTATCAT
GTCTGTATACCGTCGACCTCTAGCTAGAGCTTGGCGTAATCATGGTCATAGCTGTTTCCTGTGTGAAAT
TGTTATCCGCTCACAATTCCACACAACATACGAGCCGGAAGCATAAAGTGTAAAGCCTGGGGTGCCTA
ATGAGTGAGCTAACTCACATTAATTGCGTTGCGCTCACTGCCCGCTTTCCAGTCGGGAAACCTGTCGT
GCCAGCTGCATTAATGAATCGGCCAACGCGCGGGGAGAGGCGGTTTGCGTATTGGGCGCTCTTCCG
CTTCCTCGCTCACTGACTCGCTGCGCTCGGTCGTTCGGCTGCGGCGAGCGGTATCAGCTCACTCAA
AGGCGGTAATACGGTTATCCACAGAATCAGGGGATAACGCAGGAAAGAACATGTGAGCAAAAGGCCA
GCAAAAGGCCAGGAACCGTAAAAAGGCCGCGTTGCTGGCGTTTTTCCATAGGCTCCGCCCCCCTGA
CGAGCATCACAAAAATCGACGCTCAAGTCAGAGGTGGCGAAACCCGACAGGACTATAAAGATACCAG
GCGTTTCCCCCTGGAAGCTCCCTCGTGCGCTCTCCTGTTCCGACCCTGCCGCTTACCGGATACCTGT
CCGCCTTTCTCCCTTCGGGAAGCGTGGCGCTTTCTCATAGCTCACGCTGTAGGTATCTCAGTTCGGT
GTAGGTCGTTCGCTCCAAGCTGGGCTGTGTGCACGAACCCCCCGTTCAGCCCGACCGCTGCGCCTT
ATCCGGTAACTATCGTCTTGAGTCCAACCCGGTAAGACACGACTTATCGCCACTGGCAGCAGCCACT
GGTAACAGGATTAGCAGAGCGAGGTATGTAGGCGGTGCTACAGAGTTCTTGAAGTGGTGGCCTAACT
ACGGCTACACTAGAAGAACAGTATTTGGTATCTGCGCTCTGCTGAAGCCAGTTACCTTCGGAAAAAGA
GTTGGTAGCTCTTGATCCGGCAAACAAACCACCGCTGGTAGCGGTGGTTTTTTTGTTTGCAAGCAGC
AGATTACGCGCAGAAAAAAAGGATCTCAAGAAGATCCTTTGATCTTTTCTACGGGGTCTGACGCTCAG
TGGAACGAAAACTCACGTTAAGGGATTTTGGTCATGAGATTATCAAAAAGGATCTTCACCTAGATCCTT
TTAAATTAAAAATGAAGTTTTAAATCAATCTAAAGTATATATGAGTAAACTTGGTCTGACAGTTACCAATG
CTTAATCAGTGAGGCACCTATCTCAGCGATCTGTCTATTTCGTTCATCCATAGTTGCCTGACTCCCCGT
CGTGTAGATAACTACGATACGGGAGGGCTTACCATCTGGCCCCAGTGCTGCAATGATACCGCGAGAC
CCACGCTCACCGGCTCCAGATTTATCAGCAATAAACCAGCCAGCCGGAAGGGCCGAGCGCAGAAGT
GGTCCTGCAACTTTATCCGCCTCCATCCAGTCTATTAATTGTTGCCGGGAAGCTAGAGTAAGTAGTTC
GCCAGTTAATAGTTTGCGCAACGTTGTTGCCATTGCTACAGGCATCGTGGTGTCACGCTCGTCGTTTG
GTATGGCTTCATTCAGCTCCGGTTCCCAACGATCAAGGCGAGTTACATGATCCCCCATGTTGTGCAAA
AAAGCGGTTAGCTCCTTCGGTCCTCCGATCGTTGTCAGAAGTAAGTTGGCCGCAGTGTTATCACTCAT
GGTTATGGCAGCACTGCATAATTCTCTTACTGTCATGCCATCCGTAAGATGCTTTTCTGTGACTGGTGA
GTACTCAACCAAGTCATTCTGAGAATAGTGTATGCGGCGACCGAGTTGCTCTTGCCCGGCGTCAATAC
GGGATAATACCGCGCCACATAGCAGAACTTTAAAAGTGCTCATCATTGGAAAACGTTCTTCGGGGCGA
AAACTCTCAAGGATCTTACCGCTGTTGAGATCCAGTTCGATGTAACCCACTCGTGCACCCAACTGATC
TTCAGCATCTTTTACTTTCACCAGCGTTTCTGGGTGAGCAAAAACAGGAAGGCAAAATGCCGCAAAAA
AGGGAATAAGGGCGACACGGAAATGTTGAATACTCATACTCTTCCTTTTTCAATATTATTGAAGCATTTA
TCAGGGTTATTGTCTCATGAGCGGATACATATTTGAATGTATTTAGAAAAATAAACAAATAGGGGTTCCG
CGCACATTTCCCCGAAAAGTGCCACCTGACGTC
